# N-terminal mutant Huntingtin deposition correlates with CAG repeat length and disease onset, but not neuronal loss in Huntington’s disease

**DOI:** 10.1101/2022.05.03.490349

**Authors:** Florence E. Layburn, Adelie Y. S. Tan, Nasim F. Mehrabi, Maurice A. Curtis, Lynette J. Tippett, Nathan Riguet, Lorène Aeschbach, Hilal A. Lashuel, Mike Dragunow, Richard L. M. Faull, Malvindar K. Singh-Bains

**Affiliations:** Centre for Brain Research, University of Auckland, Private Bag 92019, Auckland 1142, New Zealand; Department of Anatomy and Medical Imaging, University of Auckland, Private Bag 92019, Auckland 1142, New Zealand; Department of Pharmacology and Clinical Pharmacology, University of Auckland, Private Bag 92019, Auckland 1142, New Zealand; Department of Psychology, University of Auckland, Private Bag 92019, Auckland 1142, New Zealand; Laboratory of Molecular and Chemical Biology of Neurodegeneration, Brain Mind Institute, Ecole Polytechnique Fédérale de Lausanne (EPFL), 1015, Lausanne, Switzerland

**Keywords:** Huntington’s disease, Middle Temporal Gyrus, Cortex, Human Brain, Tissue Microarrays, Immunohistochemistry

## Abstract

Huntington’s disease (HD) is caused by a CAG repeat expansion mutation in the gene encoding the huntingtin (Htt) protein, with mutant Htt protein subsequently forming aggregates within the brain. Mutant Htt is a current target for novel therapeutic strategies for HD, however, the lack of translation from preclinical research to disease-modifying treatments highlights the need to improve our understanding of the role of Htt protein in the human brain. This study aims to undertake a high-throughput screen of 12 candidate antibodies against various sequences along the Htt protein to characterize Htt distribution and expression in post-mortem human brain tissue microarrays (TMAs).

Immunohistochemistry was performed on middle temporal gyrus TMAs comprising of up to 28 HD and 27 age-matched control cases, using 12 antibodies specific to various sequences along the Htt protein. From this study, six antibodies directed to the Htt N-terminus successfully immunolabelled human brain tissue. The Htt aggregates and Htt protein expression levels for the six successful antibodies were subsequently quantified with high-throughput analysis. Htt aggregates were detected in HD cases using antibodies MAB5374, MW1, and EPR5526, despite no change in overall Htt protein expression compared to control cases, suggesting a redistribution of Htt into aggregates in HD. Significant associations were found between the number of Htt aggregates and both age of disease onset, and CAG repeat length in HD. However, the number of Htt aggregates did not correlate with the degree of striatal degeneration or the degree of cortical neuron loss. Together, these results suggest that longer CAG repeat lengths correlate with Htt aggregation in the HD human brain, and Htt cortical aggregate deposition is associated with the onset of clinical symptoms. This study also reinforces that antibodies MAB5492, MW8, and 2B7 which have been utilized to characterize Htt in animal models of HD are not specific for Htt in human brain tissue, thereby highlighting the need for validated means of Htt detection to support drug development for HD.

## Introduction

Huntington’s disease (HD) is a hereditary neurodegenerative disease characterized by a triad of motor, mood, and cognitive symptoms (Martin & Gusella, 1986). HD is caused by an autosomal dominantly inherited cytosine-adenine-guanine (CAG) trinucleotide repeat expansion in the IT15 gene on chromosome 4, encoding the huntingtin (Htt) protein (The Huntington’s Disease Collaborative Research Group, 1993). The formation of mutant Htt protein aggregates within the brain is hypothesized to trigger pathogenic processes causing progressive neuronal cell death, primarily affecting the basal ganglia and cerebral cortex (Herndon et al., 2009; Kim et al., 2014; Mehrabi et al., 2016; Nana et al., 2014; Singh-Bains et al., 2016; Thu et al., 2010; Tippett et al., 2007). Most therapies currently being developed for HD focus on lowering Htt production, either by lowering all Htt expression or specifically lowering mutant Htt, however, no clinical trials have achieved sufficient safety or succeeded at modifying HD progression (Mullard, 2019; Tabrizi et al., 2019). There is still controversy over the relative contributions of both normal and mutant Htt in HD, therefore developing effective tools to study Htt protein within the human brain is essential to inform development of disease-modifying treatments. The failure of clinical trials may signal the detrimental effects of targeting the entire Htt protein by inhibiting the non-pathogenic, physiologically relevant functions of Htt in the HD brain. Therefore, a thorough investigation of Htt distribution, expression, and localization within the human brain is warranted to determine the true implications of targeting Htt to treat HD.

The striatum and globus pallidus are major sites of neuronal cell loss in HD, and as such, pathology in the basal ganglia forms the basis of the neuropathological grading system for classifying HD disease severity (Vonsattel et al., 1985). The cortex is another key site of neurodegeneration in HD, with pyramidal cell loss observed in numerous cortical regions including the premotor cortex (involved in planning movement), motor cortex (involved in generating movement), somatosensory cortex (sensory information processing), cingulate gyrus (involved in mood) and the temporal cortex (involved in cognition and memory) (Halliday et al., 1998; Nana et al., 2014; Thu et al., 2010). Several anatomical studies from our laboratory have demonstrated the significant contribution of cortical neuronal cell loss towards symptom phenotype and severity in HD post-mortem cases with different symptom profiles (Kim et al., 2014; Mehrabi et al., 2016; Nana et al., 2014; Thu et al., 2010). For example, greater neurodegeneration in the motor cortex is observed in cases with predominantly motor symptoms, whereas little to no cell loss is seen in this region in cases with predominantly mood symptoms (Kim et al., 2014; Mehrabi et al., 2016; Thu et al., 2010). These relationships reinforce HD to be a complex disorder involving both subcortical and cortical structures, and the lesser degree of neuronal loss in the cerebral cortex relative to the striatum may provide insight into the early pathogenic events triggering cell death in HD (Halliday et al., 1998). Although the exact mechanism driving cell death in HD is unknown, the trigger may lie in the formation of intracellular mutant Htt protein aggregates.

Early immunohistochemical studies of the post-mortem HD brain have demonstrated greater Htt aggregate deposition throughout the cerebral cortex relative to the striatum (Becher et al., 1998; DiFiglia et al., 1997; Gutekunst et al., 1999; Halliday et al., 1998; Sapp et al., 1997). However, the contribution of Htt aggregates to HD pathogenesis has not yet been fully elucidated. Evidence from previous studies aiming to map Htt deposition in the post-mortem human HD brain suggests a key role for mutant Htt N-terminal fragments in HD pathology, for example, aggregates could only be identified by immunolabelling the N-terminus of the protein (Becher et al., 1998; DiFiglia et al., 1997; Sapp et al., 1997; Sharp et al., 1995). Subsequent mechanistic studies utilizing *in vitro* models of HD demonstrated that mutant Htt undergoes cleavage of its polyglutamine (polyQ)-containing N-terminus, with accumulation of the resulting fragments as insoluble aggregates within the cytoplasm and nuclei of neurons (Cooper et al., 1998; Schilling et al., 2007). Furthermore, transgenic mice that express the mutant Htt N-terminus display neurotoxicity and Htt aggregate formation that closely resembles what is observed in the human HD brain (Davies et al., 1997; O’Kusky et al., 1999).

The most commonly used antibodies to profile Htt aggregates (1C2, 1F8, S830, and EM48) in human brain immunohistochemical studies bind to Htt at its N-terminus (Gutekunst et al., 1999; Herndon et al., 2009). Although 1C2, 1F8, S830, and EM48 have served as valuable tools for profiling Htt aggregates, each one has limitations. For example, 1C2 and 1F8 were both raised against the TATA binding protein and are not strictly specific for human Htt, while S830 is not commercially available (Trottier et al., 1995; White et al., 1997). It should also be noted that the majority of studies to map Htt aggregates have been carried out in HD mouse models of HD, whereby Htt aggregation is induced by a non-physiological CAG repeat length expansion, with each mouse line demonstrating a unique Htt immunoreactivity profile (Bayram-Weston et al., 2016). Furthermore, mouse models of HD do not recapitulate the diversity of Htt fragments in the human HD brain in which multiple types of Htt N-terminal fragments have been observed, with differences in fragment length giving rise to distinct structures and mechanisms of aggregation (Kolla et al., 2021). Given the biochemical and structural diversity of Htt protein in the human HD brain, it is unlikely that a single antibody is capable of capturing all pathogenic Htt protein, highlighting the need to develop antibodies which (1) are specific to human Htt protein, (2) represent sequences along the entire length of Htt protein, and (3) can accurately distinguish between the normal and mutant forms.

This study aimed to provide a comprehensive profile of Htt immunoreactivity in both neurologically normal control and HD human brain samples by screening cortical brain tissue microarrays (TMAs). To address this aim, immunohistochemistry was carried out using 12 antibodies specific to various amino acid sequences along the Htt protein, as illustrated in Figure 1 A. Furthermore, the relationship between Htt expression identified with successful candidate antibodies was compared with the degree of neuronal cell loss and other HD clinico-pathological characteristics such as CAG repeat length, Vonsattel neuropathological grade, and age of disease onset. Based on the current body of literature, we hypothesized that more Htt aggregates will be detected with antibodies directed against the N-terminus compared to the C-terminus of Htt, and that more Htt aggregates will be observed in HD compared to control cases. However, considering the results of previous anatomical studies of the HD brain, we did not predict a relationship between Htt aggregate distribution and the extent of neuronal cell loss in the HD cases. We hypothesized that the application of multiple antibodies targeting different Htt sequences will increase our chances of capturing different types of Htt pathological aggregates, due to differences in both their structural and biochemical properties, which may form at different conditions, in different cell types, and from different Htt fragments. Furthermore, the application of Htt antibodies to human brain TMAs with up to 55 human tissue samples will capture HD case heterogeneity, which is essential for understanding a disease with variable clinicopathology.

**Figure 1.**
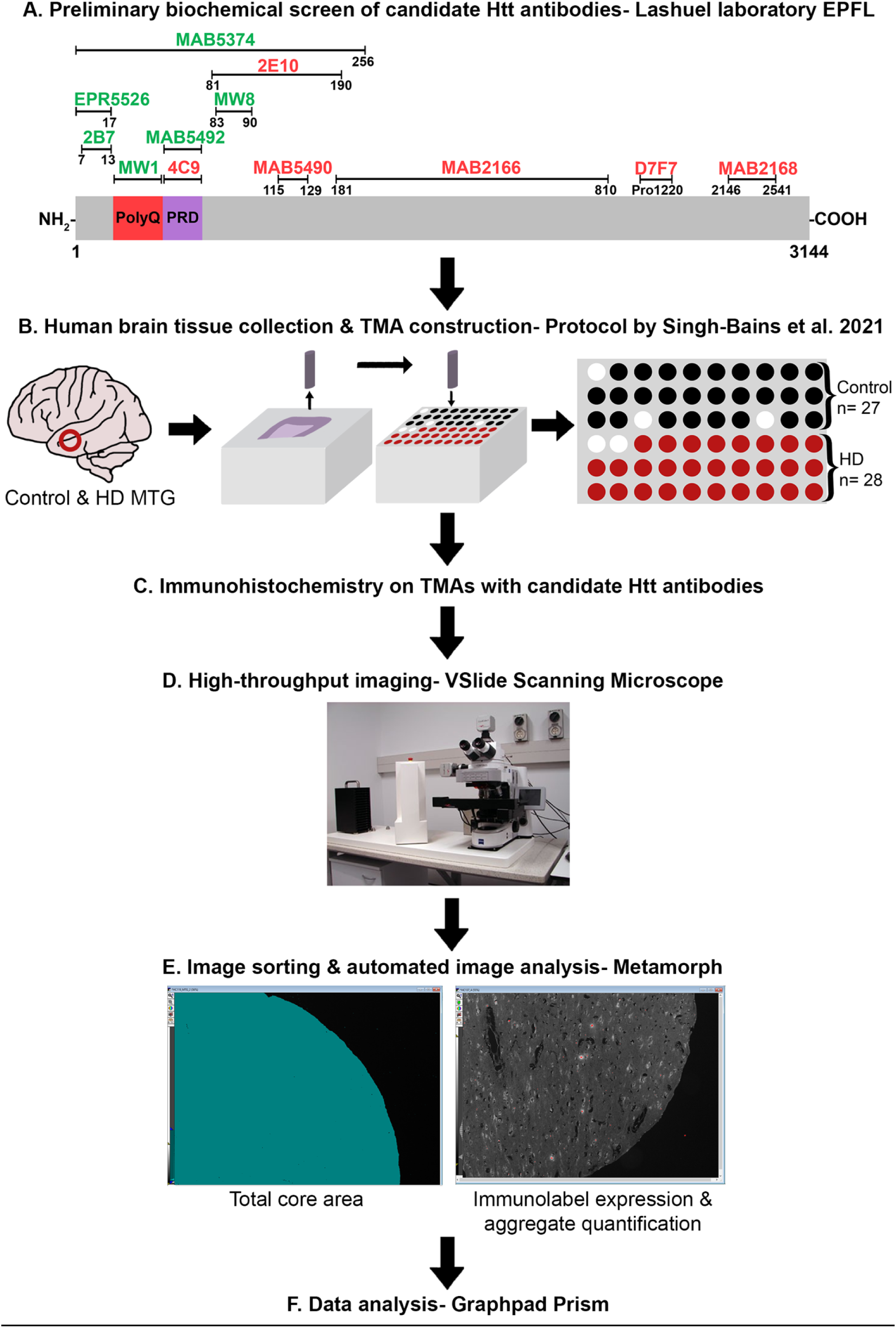
Workflow of Htt antibody screening on human brain tissue microarrays (TMAs). (A) A range of primary anti-Htt antibodies were biochemically screened by the Lashuel laboratory, Ecole Polytechnique Fédérale de Lausanne. 12 antibodies binding various regions of Htt protein were selected and supplied for immunohistochemical screening in this study. (B) MTG human brain tissue was collected and utilized to construct TMAs according to the protocol by Singh-Bains et al., (2021). The TMA map schematic denotes the 28 HD and 27 neurologically normal control human brain tissue samples included in each TMA. (C) Immunohistochemistry was performed with candidate anti-Htt antibodies on the TMAs for subsequent quantitative studies. (D) High-throughput automated image acquisition was performed using the VSlide Scanning Microscope. (E) The images from (D) were sorted to exclude missing and damaged TMA cores, then analyzed using Metamorph software to determine the total core area, immunolabel expression, and aggregate number detected using each anti-Htt antibody. (F) Data analysis for comparisons between the control and HD cohorts was performed using GraphPad Prism. Htt = huntingtin; TMA = tissue microarray; MTG = middle temporal gyrus; HD = Huntington’s disease.

## Materials and Methods

### Human brain tissue collection and preparation

Post-mortem human brain tissue utilized in this project was obtained from the Neurological Foundation Human Brain Bank in the Centre for Brain Research, the University of Auckland. All research procedures and protocols utilizing human brain tissue were conducted under full ethics approval from the Health and Disability Ethics Committee (Ref.14/NTA/208) with informed consent from both families and donors.

Fixation and dissection protocols for human brain tissue preparation have been described previously (Waldvogel et al., 2006; Waldvogel et al., 2008). Briefly, the brains were perfused and, for each case, a small ∼3 mm thick portion of the middle temporal gyrus (MTG) was embedded in paraffin wax to be utilized for the construction of tissue microarrays (TMAs). The MTG has previously been implicated in HD pathogenesis, with a demonstrated reduction in SMI-32^+^ pyramidal neurons in HD compared to the normal control brains, confirming neuronal cell loss and thereby justifying the use of this region for further HD neuropathology studies (Nana et al., 2014). The TMAs consist of post-mortem brain tissue from the MTG of 27 neurologically normal control cases and 28 Huntington’s disease (HD) cases (Table 1 and Table 2). The control and HD cases were matched as closely as possible for age at death, sex, and post-mortem delay (PMD).

**Table 1.**
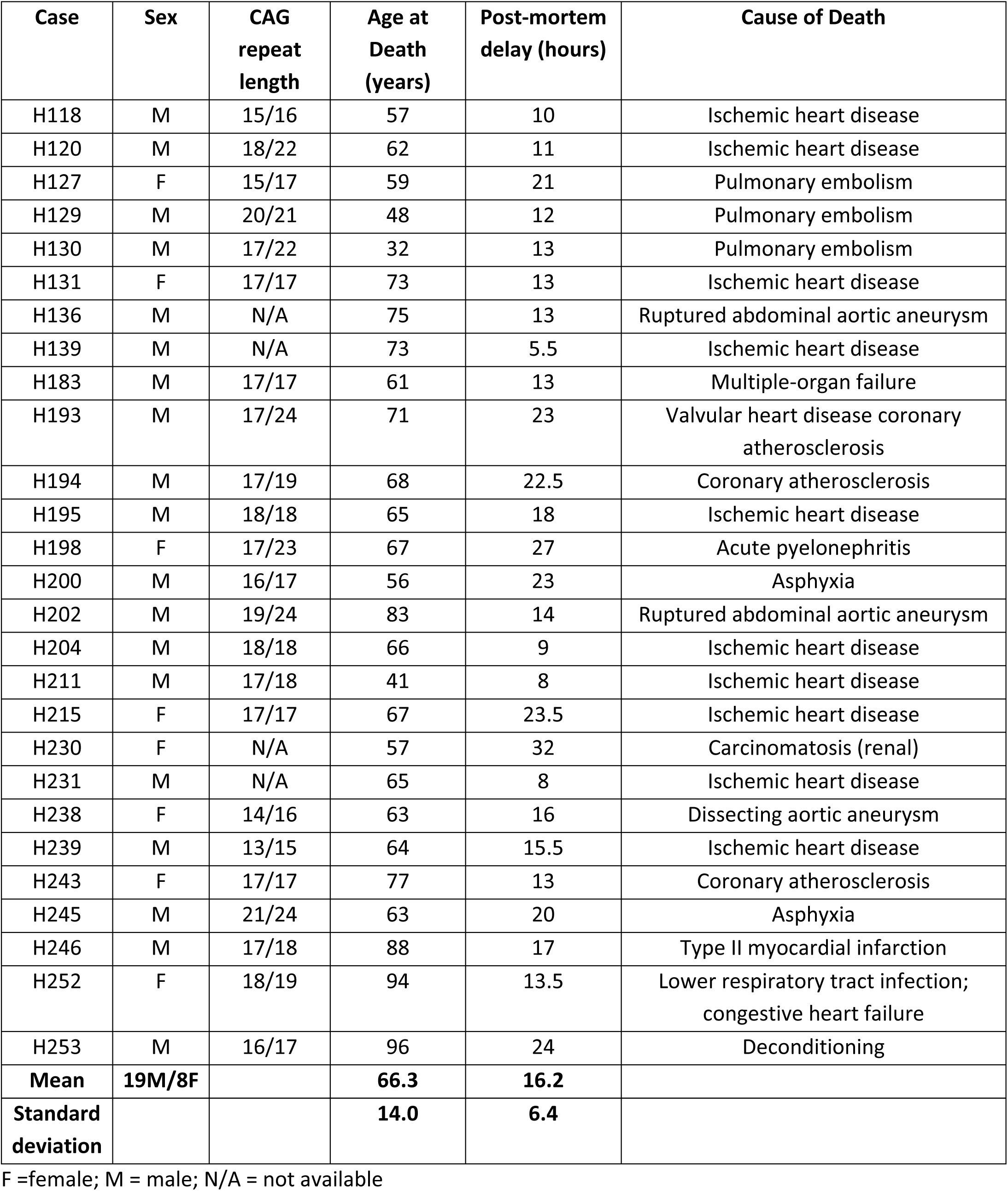
Neurologically normal control cases for cortical tissue microarray (TMA 18C)

| Case | Sex | CAG repeat length | Age at Death (years) | Post-mortem delay (hours) | Cause of Death |
| --- | --- | --- | --- | --- | --- |
| H118 | M | 15/16 | 57 | 10 | Ischemic heart disease |
| H120 | M | 18/22 | 62 | 11 | Ischemic heart disease |
| H127 | F | 15/17 | 59 | 21 | Pulmonary embolism |
| H129 | M | 20/21 | 48 | 12 | Pulmonary embolism |
| H130 | M | 17/22 | 32 | 13 | Pulmonary embolism |
| H131 | F | 17/17 | 73 | 13 | Ischemic heart disease |
| H136 | M | N/A | 75 | 13 | Ruptured abdominal aortic aneurysm |
| H139 | M | N/A | 73 | 5.5 | Ischemic heart disease |
| H183 | M | 17/17 | 61 | 13 | Multiple-organ failure |
| H193 | M | 17/24 | 71 | 23 | Valvular heart disease coronary atherosclerosis |
| H194 | M | 17/19 | 68 | 22.5 | Coronary atherosclerosis |
| H195 | M | 18/18 | 65 | 18 | Ischemic heart disease |
| H198 | F | 17/23 | 67 | 27 | Acute pyelonephritis |
| H200 | M | 16/17 | 56 | 23 | Asphyxia |
| H202 | M | 19/24 | 83 | 14 | Ruptured abdominal aortic aneurysm |
| H204 | M | 18/18 | 66 | 9 | Ischemic heart disease |
| H211 | M | 17/18 | 41 | 8 | Ischemic heart disease |
| H215 | F | 17/17 | 67 | 23.5 | Ischemic heart disease |
| H230 | F | N/A | 57 | 32 | Carcinomatosis (renal) |
| H231 | M | N/A | 65 | 8 | Ischemic heart disease |
| H238 | F | 14/16 | 63 | 16 | Dissecting aortic aneurysm |
| H239 | M | 13/15 | 64 | 15.5 | Ischemic heart disease |
| H243 | F | 17/17 | 77 | 13 | Coronary atherosclerosis |
| H245 | M | 21/24 | 63 | 20 | Asphyxia |
| H246 | M | 17/18 | 88 | 17 | Type II myocardial infarction |
| H252 | F | 18/19 | 94 | 13.5 | Lower respiratory tract infection; congestive heart failure |
| H253 | M | 16/17 | 96 | 24 | Deconditioning |
| **Mean** | **19M/8F** |  | **66.3** | **16.2** |  |
| **Standard deviation** |  |  | **14.0** | **6.4** |  |

F =female; M = male; N/A = not available

**Table 2.** Huntington’s disease cases for the cortical tissue microarray (TMA 18C)

| Case | Sex | CAG repeat length | Age of Death (years) | PMD (hours) | Age of Onset (years) | Vonsattel Striatal Neuropathological Grade | Cause of Death |
| --- | --- | --- | --- | --- | --- | --- | --- |
| HC051 | M | 16/43 | 58 | 4.5 | 40 | 1 | Pneumonia |
| HC052 | F | 19/46 | 61 | 23 | 31 | 2 | Cerebro-vascular accident |
| HC075 | M | 18/52 | 41 | 5.5 | 21 | 3 | Pneumonia |
| HC076 | M | 19/42 | 71 | 16 | 52 | 2 | Pneumonia |
| HC088 | F | 17/44 | 57 | 19 | 44 | 2 | Myocardial infarction |
| HC089 | F | 17/41 | 71 | 12 | 64 | 2 | Pneumonia |
| HC091 | F | 23/44 | 53 | 5.5 | 31 | 3 | HD |
| HC096 | F | 22/53 | 39 | 20 | 15 | 3 | Starvation with dehydration |
| HC102 | M | 17/42 | 64 | 10 | 33 | 3 | Aspiration pneumonia |
| HC104 | M | 18/51 | 40 | 15 | 15 | 3 | Pneumonia |
| HC119 | M | 17/48 | 51 | 15.5 | 31 | 3 | Dehydration |
| HC123 | M | 23/47 | 54 | 14 | 31 | 4 | Bronchopneumonia |
| HC128 | M | N/A | 80 | 13 | 50 | 3 | Myocardial infarction |
| HC133 | M | 17/43 | 65 | 13 | 48 | 2 | Renal failure |
| HC134 | M | 18/43 | 62 | 9 | 40 | 2 | HD |
| HC137 | M | 17/41 | 83 | 13 | 40 | 1 | Pneumonia |
| HC140 | M | 17/40 | 62 | 22 | 44 | 3 | Chronic renal failure |
| HC141 | F | -/42 | 84 | 14 | 70 | 3 | N/A |
| HC142 | F | 15/45 | 55 | 25 | 35 | 3 | Pneumonia |
| HC143 | F | 17/54 | 45 | 16 | 21 | 4 | Respiratory arrest |
| HC144 | M | 10/46 | 55 | N/A | 44 | 1 | Leukaemia |
| HC145 | M | N/A | 54 | N/A | 41 | 3 | Chest infection |
| HC146 | M | 17/47 | 46 | 6 | 30 | 3 | HD |
| HC147 | M | 27/42 | 64 | 18 | 30 | 3 | Pulmonary thromboembolus |
| HC148 | M | 22/43 | 63 | 16 | 39 | 3 | Pulmonary embolism |
| HC150 | F | 22/42 | 64 | 21 | 51 | 2 | Bronchopneumonia |
| HC151 | M | 17/43 | 71 | 35.5 | 48 | 3 | Pre-renal failure, poor fluid intake and swallowing |
| HC152 | M | 23/43 | 60 | 16 | 38 | 2 | Renal failure |
| **Mean** | **18M/10F** |  | **59.7** | **15.3** | **38.5** |  |  |
| **Standard deviation** |  |  | **11.7** | **6.7** | **12.7** |  |  |

F =female; M = male; N/A = not available

The control cases included in the TMAs (Table 1) consisted of 19 males and 8 females, with an age at death of 32-96 years [mean = 66.3 ± 14.0 years old (mean ± SD)], and PMD of 8-24 hours (mean = 16.2 ± 6.4 hours). The HD cases included in the TMAs consisted of 18 males and 10 females, with an age at death of 39-84 years (mean = 59.7 ± 11.7 years old), and PMD of 4.5-35.5 hours (mean = 15.3 ± 6.7 hours). The age of disease onset for HD cases was 15-70 years (mean = 38.5 ± 12.7 years old). Age of clinical onset was based on well-defined criteria outlined by Tippet et al., (2007). For both normal control and HD cases, the polyglutamine (CAG) repeat length was determined using polymerase chain reaction from either a blood sample or cerebellar brain tissue sample (Table 1 and 2). HD cases (Table 1) were neuropathologically graded based on the degree of striatal atrophy according to the Vonsattel grading system (Vonsattel et al., 1985).

### Tissue microarray (TMA) construction

The human brain TMAs utilized in this study were constructed according to the protocol by Singh-Bains et al. (2021). The TMA utilized in this study, designated as TMA 18C, consists of cylindrical cores of formalin-fixed, paraffin-embedded human brain tissue obtained from the MTG of 27 neurologically normal control and 28 HD cases (Table 1 and Table 2). Prior to TMA construction, a section from each designated donor block of human brain tissue was stained with cresyl violet (Nissl stain) to enable the identification of layers II-V of the cerebral cortex and provide a guide for tissue coring, ensuring only grey matter was captured for each case. An adjacent section was immunolabelled with ubiquitously expressed antibody anti-glial fibrillary acidic protein (GFAP) to ensure antigenicity of HD and control tissue for TMA utilization. Once the neuroanatomical and antigenicity studies were complete, 2mm-diameter cores were extracted from each donor block of tissue and inserted in a recipient block of paraffin wax using the Veridium VTA-100 Tissue Microarrayer. Once the coring process was complete, the TMA recipient block was cut into 7μm sections using a microtome (RM2235, Leica) and mounted onto positively charged slides (Uber Frost) in preparation for immunohistochemistry.

### Tissue microarray immunohistochemistry

Immunohistochemistry (IHC) was conducted using 12 monoclonal antibodies targeting different amino acid sequences of the Huntingtin (Htt) protein, as summarized in Table 3. Standard 3,3’-diaminobenzidine (DAB) IHC protocols were applied, as previously described (Singh-Bains et al., 2021). Briefly, the TMA sections were placed on a hotplate to anneal the tissue to the slides (1 hour at 60°C), dewaxed in xylene (twice; for 1 hour and then 10 min), rehydrated through an ethanol series (100%, 95%, 80%, 75%), and then washed in milliQ water (mQH_2_O) (3 × 5 min). Next, the slides were transferred into antigen retrieval buffer for 20 minutes at 121°C (model 2100-retriever, Pick Cell Laboratories) (Additional file: Supplementary Table 1). The slides were subsequently left to cool at room temperature for 2 hours and washed in mQH_2_O (3 × 5 min). As an additional antigen retrieval step, the slides were submerged in 100% formic acid for 5 minutes, before washing again in mQH_2_O (3 × 5 min).

**Table 3.** All primary monoclonal anti-Huntingtin antibodies trialled for immunohistochemistry in this study. All primary antibodies used in this study were provided by Professor Hilal Lashuel, École Polytechnique Fédérale de Lausanne.

| **Antibody** | **Huntingtin specificity**<br>**(amino acid sequence)** | **Antigen used in production** | **Host species** | **Company** | **Category number** | **Lot number/**<br>**Clone number** |
| --- | --- | --- | --- | --- | --- | --- |
| **2B7** | N-terminal 7-13 | Human Htt | Mouse | Thermo Scientific | CHDI-90000830-2 | Clone 2B7 |
| **EPR5526** | N-terminal amino acids 1-17 | Synthetic peptide | Rabbit | Abcam | ab109115 | Lot GR3208603-2 |
| **MAB5374** | N-terminal 1-256 with the deletion of the polyQ tract | GST fusion protein | Mouse | Millipore | MAB5374 | Lot 3052503 |
| **MAB5492** | Proline-rich domain | Recombinant human Htt | Mouse | Millipore | MAB5492 | Lot 3086659 |
| **MW1** | Expanded polyQ tract | GST tagged recombinant protein | Mouse | Thermo Scientific | CHDI-90000895-5 | Clone 1G101G12 |
| **MW8** | N-terminal 83-90 | Human Htt exon 1 with 67 polyQ aggregate | Mouse | Thermo Scientific | CHDI-90000942-1 | Clone 18C52B3 |
| **2E10** | N-terminal 81-190 | GST tagged partial recombinant protein | Mouse | Abnova | H00003064-M01 | Lot J5151-2E10 |
| **4C9** | N-terminal 51-71; proline-rich domain | Human Htt | Mouse | Thermo Scientific | CHDI-90000833-3 | Clone 4C9 |
| **D7F7** | Residues surrounding Pro1220 | Synthetic peptide | Rabbit | Cell Signalling | 5656S | Lot 4 |
| **MAB2166** | Internal segment 181-810 | Fusion protein | Mouse | Millipore | MAB2166 | Lot 3068437 |
| **MAB2168** | C-terminal 2146-2541 | Fusion protein | Mouse | Millipore | MAB2168 | Lot 2907694 |
| **MAB5490** | N-terminal 115-129 | Recombinant human Htt | Mouse | Millipore | MAB5490 | Lot 3184135 |

Next, an endogenous peroxidase block was carried out by immersing the slides in a blocking solution of 50% methanol and 1% H_2_O_2_, diluted in mQH_2_O (20 min). The slides were rinsed in mQH_2_O, then washed in phosphate-buffered saline (PBS) (2 × 5 min). The slides were incubated with 10% normal goat serum (diluted in PBS) for 1 hour at room temperature. Next, the slides were incubated with the primary antibody at 4°C overnight (Table 3).

The following day, the slides were washed with PBS with 0.1% Triton X-100 (PBS-T) for 5 minutes, then with PBS (2 × 5 min) to wash off unbound primary antibody. The slides were incubated with the secondary antibodies, either goat anti-mouse (1:500; Sigma Aldrich; Cat #B7264-2ML; Lot #SLBK5415V) or goat anti-rabbit (1:500 Sigma Aldrich; Cat #B7389-2ML; Lot #SLBR0916V), at room temperature for 3 hours. The slides were washed (5 min with PBS-T; 2 × 5 min with PBS), then incubated with ExtrAvidin® Peroxidase (1:1000 Sigma Aldrich; Cat #E2886-1ML; Lot #047M4805V) at room temperature for 1 hour. The slides were washed in PBS-T and PBS again, before incubation with DAB solution with 0.04% nickel ammonium sulphate for a period of 5-15 minutes (Additional file: Supplementary Table 1). Once the incubation with DAB was complete, the slides were washed in PBS (3 × 5 min), followed by mQH_2_O (3 × 5 min). The slides were subsequently dehydrated by submersion in ethanol (75%, 80%, 95%, 100%) and xylene (3 × 10 min). Finally, the slides were coverslipped using DPX as a mounting agent and left to dry overnight before imaging. A no-primary control was included in each experiment to assess the degree of non-specific background staining, with 1% normal goat serum applied instead of the primary antibody. All primary antibodies used in this study were first optimised for IHC in whole paraffin-embedded MTG tissue sections from both neurologically normal control and HD human brains. The optimized concentrations and experimental conditions required for each antibody are summarized in Supplementary Table 1.

### Tissue microarray image acquisition and sorting

The image acquisition procedures utilized in this study are outlined in the protocol by Singh-Bains et al., (2021). Once the TMAs were successfully immunolabelled, images of the TMA cores were acquired using the V-Slide Automated Slide Scanning Microscope (Metasystems) and Metafer5 software (version 3.16.160) at the Biomedical Imaging Research Unit (BIRU), the University of Auckland. Briefly, the imaging protocol involved an initial pre-scan step to capture the whole TMA at low power, obtaining 75 images using a 2.5x magnification lens. The “Microarray Tool” function was then used to align a grid of 6 x 10 dots to then center of each TMA core on the pre-scan images. This step enabled the secondary re-scan step to obtain four images using a 10x magnification lens around the center of each core, acquiring a total of 240 images per TMA (4 images per core, with up to 60 cores imaged per TMA). A “stitch” image of the whole immunolabelled TMA was also generated from the pre-scan images and used in the subsequent image sorting steps. The 240 re-scan images per TMA were sorted prior to automated image analysis for the purpose of excluding missing cores and blank orientation spaces, and excluding portions of cores that were unsuitable for analysis.

### Tissue microarray image analysis

Following image acquisition and sorting, the complete TMA core re-scan images were analyzed using Metamorph software (Metamorph Offline v.7.8.10.0, Molecular Devices) according to the protocol by Singh-Bains et al. (2021). Customized Metamorph ‘journals’ were used to quantify the area of the TMA core and number of immunolabelled objects within the core images, as described below.

### Integrated Optical Density (Scaled) – Measuring total core area

The Integrated Optical Density (Scaled) journal was used to measure the total area of the immunolabeled core in pixels (Figure 2). First, the original core re-scan image was used to generate a ‘shading’ image (Figure 2 B). The ‘Optical Density (Scaled)’ function was then used to generate a 16-bit image from both the original re-scan and the ‘shading’ image, and the 16-bit image was subsequently thresholded to result in the turquoise selection shown in Figure 2 C. The ‘Thresholded Area’ (total area of the core) was exported to an Excel spreadsheet using the ‘Show Region Statistics’ function and used for calculating the immunolabel coverage for each anti-Htt marker (Figure 2 D).

**Figure 2.**
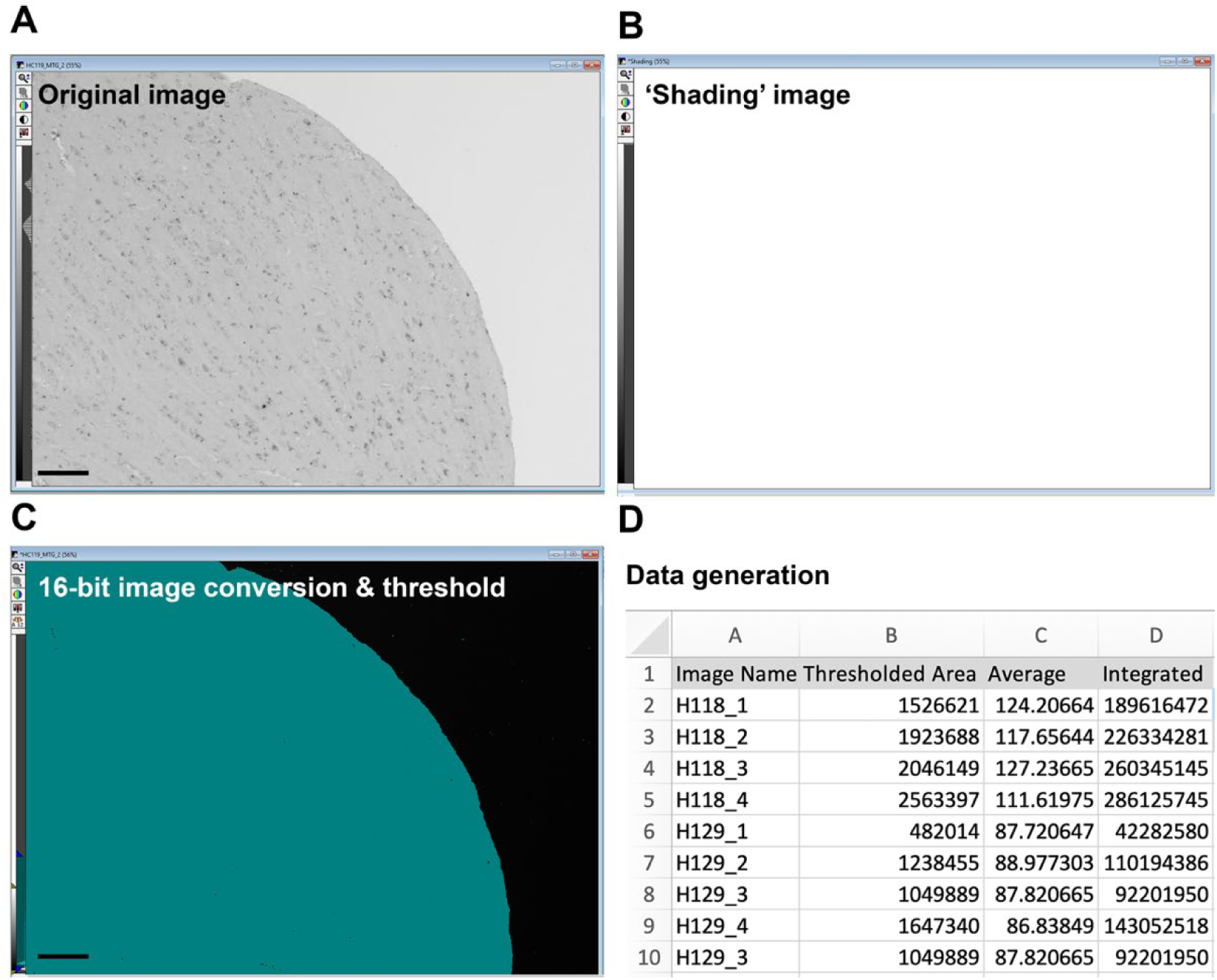
Integrated Optical Density (Scaled) journal used to measure the total area of the tissue microarray core. The (A) original re-scan core image was used to create a (B) ‘shading’ image using the ‘Add’ operation in the ‘Arithmetic’ function. The ‘Optical Density (Scaled)’ function was used to generate a 16-bit image from both the original re-scan and the ‘shading’ image (B). The ‘Threshold Image’ function was subsequently used to obtain the turquoise area (C), measuring the total area of the core image. Using the ‘Show Region Statistics’ function, the total area data were exported to an Excel spreadsheet (D). Scale bar corresponds to 100 μm.

### Count Nuclei – Measuring number and integrated intensity of objects

The Count Nuclei journal was utilised to quantify the number of Htt protein aggregates and the integrated intensity (immunolabel expression level) (Figure 3). First, the original re-scan images were inverted and converted into a 16-bit image (Figure 3 B). The ‘Count Nuclei’ function was used to identify any positive immunolabelled signals based on adjusting intensity above local background, maximum and minimum width of objects, predetermined during pilot analyses (Figure 3 C). The ‘Count Nuclei’ measurements were then exported to an Excel spreadsheet and the data for each case was expressed as a mean of four images per core.

**Figure 3.**
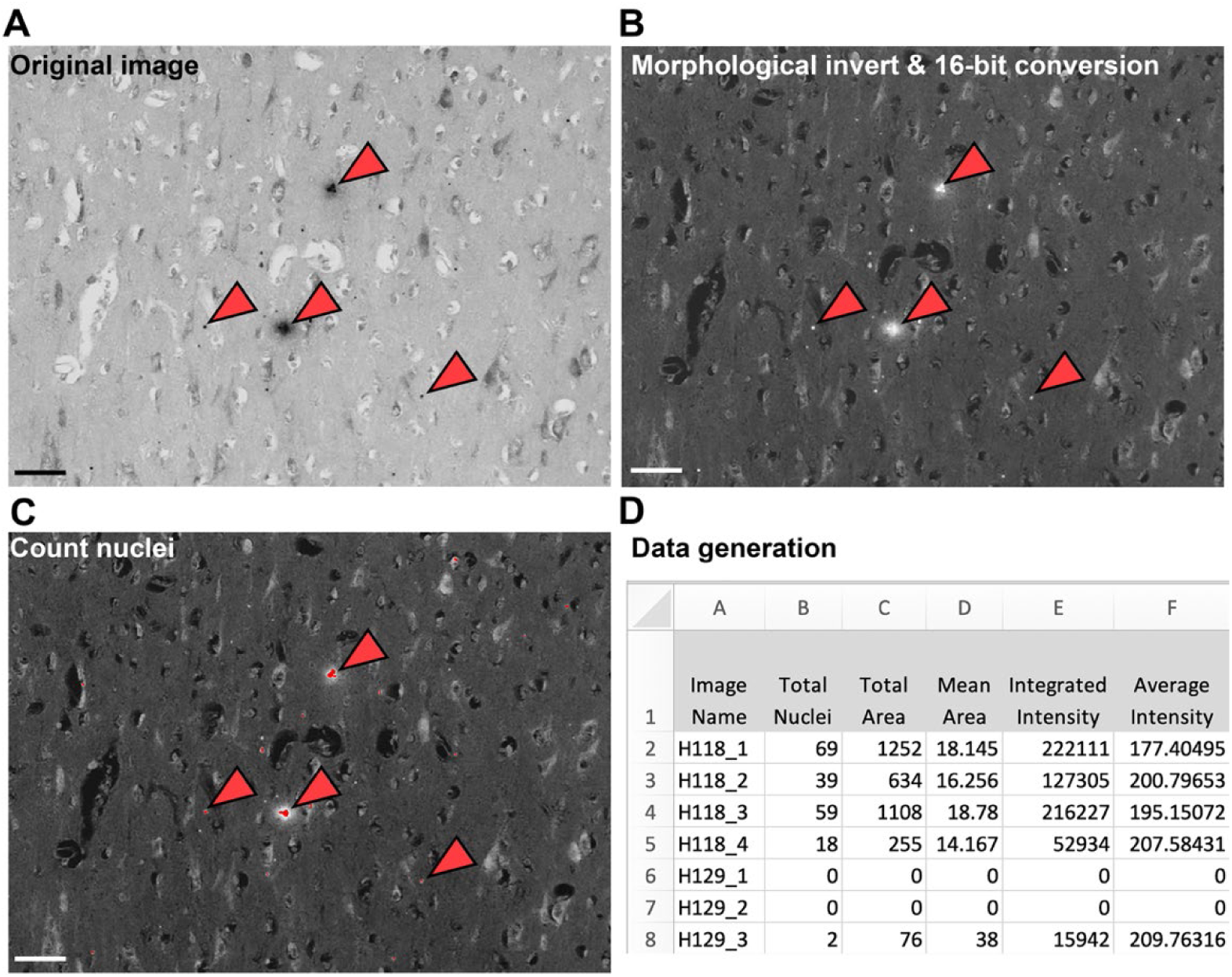
Count Nuclei journal used to measure the number of punctate objects, area, and integrated intensity (expression) of the immunolabel signal. The (A) original re-scan core image was (B) inverted using the‘Morphological Invert’function and converted into a 16-bit image using the‘Add’ operation in the‘ Arithmetic’function. Densitometric analysis was then performed on the 16-bit image using the‘Count Nuclei’ function which factored in the maximum and minimum widths of the object of interest and the intensity of the signal above local background. The objects of interest which were measured are designated in‘red’ (C), and denoted with red arrows, within the final image. The ‘Total nuclei’ and integrated intensity were exported to an Excel spreadsheet (D). Scale bar corresponds to 50 μm.

From the data generated, the ‘Total Nuclei’ measurement was used to represent the total number of Htt aggregates averaged across four images acquired per TMA core. Aggregates were classified based on the intensity of immunoreactivity above background and presence of punctate shape, which were determined using Metamorph software by thresholding immunoreactivity signals above the background levels by a factor of 150-190 and setting a maximum aggregate width of 150 pixels. ‘Total Nuclei’ data was generated for TMAs immunostained with antibodies MAB5374, MW1, EPR5526, and MAB5492 to represent the total number of Htt aggregates. The ‘Integrated Intensity’ measurement was also used to represent the total immunolabel expression levels for TMAs immunostained with MAB5374, MW1, EPR5526, MAB5492, MW8 and 2B7, capturing both aggregated and diffuse Htt immunoreactivity. ‘Integrated Intensity’ was generated from the sum of the intensity values of the pixels corresponding to the signal that was measured. Obtaining the number of Htt aggregates was unsuitable for immunostained TMAs with antibodies 2B7 and MW8, as the immunoreactivity patterns for these markers did not represent distinguishable punctate objects, therefore the ‘Integrated Intensity’ measurement was only applied.

In addition to object number and integrated intensity (immunolabel expression levels), the ‘coverage’ of each immunolabel, representing the overall coverage of the immunolabel irrespective of intensity levels, was calculated for each marker by dividing the total area of the immunolabel (generated by the Count Nuclei journal) by the total area of the core (generated by the Integrated Optical Density (Scaled) journal).

### Data analysis

Data generated from Metamorph core image analysis was statistically analysed using GraphPad Prism (GraphPad Prism 9.0.0, GraphPad Software Inc.) to compare the control and HD cohorts within the immunolabeled TMA. Data was presented as mean ± standard deviation (SD), and the parameters analysed for each marker are summarised in Table 4.

**Table 4.** Summary of parameters analysed for each anti-Huntingtin marker. The parameters were determined based on the unique pattern of immunoreactivity resulting from each marker.

| Marker | Parameters analyzed |
| --- | --- |
| MAB5374 | Aggregate number, integrated intensity, coverage |
| MW1 | Aggregate number, integrated intensity, coverage |
| EPR5526 | Aggregate number, integrated intensity, coverage |
| MAB5492 | Aggregate number, integrated intensity, coverage |
| MW8 | Integrated intensity, coverage |
| 2B7 | Integrated intensity, coverage |

First, the datasets were inspected for normality to determine the appropriate statistical test by manual inspection of a dot plot of the data in addition to conducting the D’Agostino-Pearson normality test in Prism. Based on these criteria, and the assessment of skewness, the data generated from both the control and HD cohorts was classified as not normally distributed, therefore non-parametric statistical tests were used.

For comparisons of mean aggregate numbers, mean integrated intensity, and mean immunolabel coverage between the control and HD cohorts, a non-parametric two-tailed Mann-Whitney test was carried out. The relationship between the degree of striatal neuropathology measured according to Vonsattel grade and Htt immunoreactivity was investigated for each anti-Htt marker by subgrouping the HD cases according to neuropathological grade. Mean comparisons were carried out between the control cohort, pooled HD cases of Grades 1-2, and pooled HD cases of Grades 3-4 using a Kruskal-Wallis test with Dunn’s multiple comparisons test. A non-parametric Spearman’s correlation was performed to compare SMI-32^+^ neurons (representing the degree of pyramidal neuronal loss in HD) and 1C2^+^ Htt aggregates (a surrogate marker for Htt which was raised against TATA binding protein,) respectively with the parameters investigated in this study. Correlations with a two-tailed *p*-value of < 0.05 were considered to be statistically significant, and the strength of the relationship was determined based on the Spearman’s correlation *r* coefficient as summarised in Table 5 (Taylor, 1990).

**Table 5.** Classification used for correlation strength. The Spearman’s correlation coefficient *r* value was classified according to strength. A correlation was considered statistically significant for a two-tailed *p*-value of ≤ 0.05.

| Correlation strength | r values |  |
| --- | --- | --- |
| Weak correlation | $-0.350 \leq r \leq 0$ | $0 \leq r \leq 0.350$ |
| Moderate correlation | $-0.351 \leq r \leq -0.670$ | $0.351 \leq r \leq 0.670$ |
| Strong correlation | $-0.671 \leq r \leq 0.900$ | $0.671 \leq r \leq 0.900$ |
| Very strong correlation | $r < -0.90$ | $r > 0.90$ |

Moreover, an investigation was carried out to determine if the clinical characteristics of each control and HD case were associated with Htt immunoreactivity for each anti-Htt immunolabel. A non-parametric Spearman’s correlation was performed to compare the CAG repeat length, age of disease onset, age of death, and PMD respectively with the parameters investigated in this study.

## Results

In this study, twelve anti-Htt antibodies were trialed in paraffin-embedded human brain tissue from HD and neurologically normal control cases using a variety of immunohistochemistry experimental conditions (Additional file: Supplementary Table 1). As summarized in Figure 4 and Supplementary Table 1, antibodies MAB5374, MW1, EPR5526, MAB5492, MW8, and 2B7 displayed immunoreactivity, whereas 2E10, 4C9, D7F7, MAB5490, MAB2166, and MAB2168 did not successfully immunolabel human brain tissue relative to the no-primary antibody controls. All successful immunolabels were by antibodies specific to an amino acid sequence within the N-terminus of Htt, while all C-terminal-specific antibodies did not successfully immunolabel Htt. We summarize the resultant quantitative studies carried out using the six successful antibody candidates to immunolabel HD tissue microarrays (TMAs) with a final sample size of *n* ≤ 22 control and *n* ≤ 24 HD cases.

**Figure 4.**
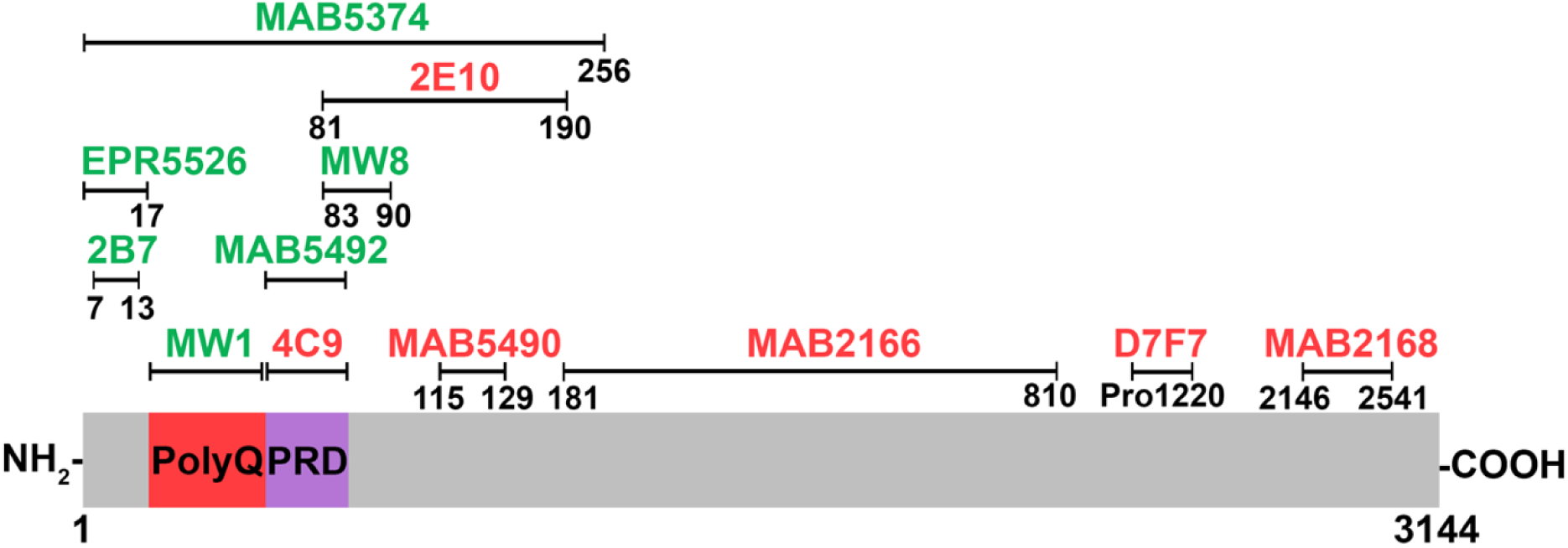
Schematic representation of candidate anti-Htt antibodies against amino acid sequences located along the Htt protein. Antibodies denoted in green successfully displayed immunoreactivity in paraffin-embedded HD and control human brain tissue immunohistochemistry. Antibodies denoted in red did not display immunoreactivity. Numbers 1 and 3144 refer to the first and last amino acid of the Htt protein sequence respectively. Htt = Huntingtin; NH_2_ = Htt N-terminus; COOH = Htt C-terminus; PolyQ = polyglutamine tract; PRD = proline-rich domain; Pro = proline; HD = Huntington’s disease.

### MAB5374 demonstrates Htt aggregate-specific immunoreactivity in HD cortical brain tissue

MAB5374 (also known as EM48) was generated against a GST-tagged fragment of the human Htt protein containing the N-terminal 256 amino acids, with the deletion of the polyglutamine (polyQ) tract (Figure 5 A) (Reindl et al., 2019). MAB5374 immunoreactivity in the TMA was specific to Htt aggregates, displaying punctate staining in HD cores only (Figure 5 B-C). High-throughput TMA analysis of MAB5374 staining demonstrated a 12-fold increase in the number of MAB5374^+^ aggregates (p<0.0001), a 16-fold increase in staining expression denoted by integrated intensity (p<0.0001), and 17-fold increase in coverage (p<0.0001) in HD cores relative to controls (Figure 5 D, F, G). Subdividing the HD cases according to Vonsattel striatal neuropathological grade (Figure 5 E) revealed no difference in MAB5374^+^ aggregates between HD Grades 1-2 and Grades 3-4, thereby confirming that the number of MAB5374^+^ aggregates is not associated with advancing striatal neuropathological grade.

**Figure 5.**
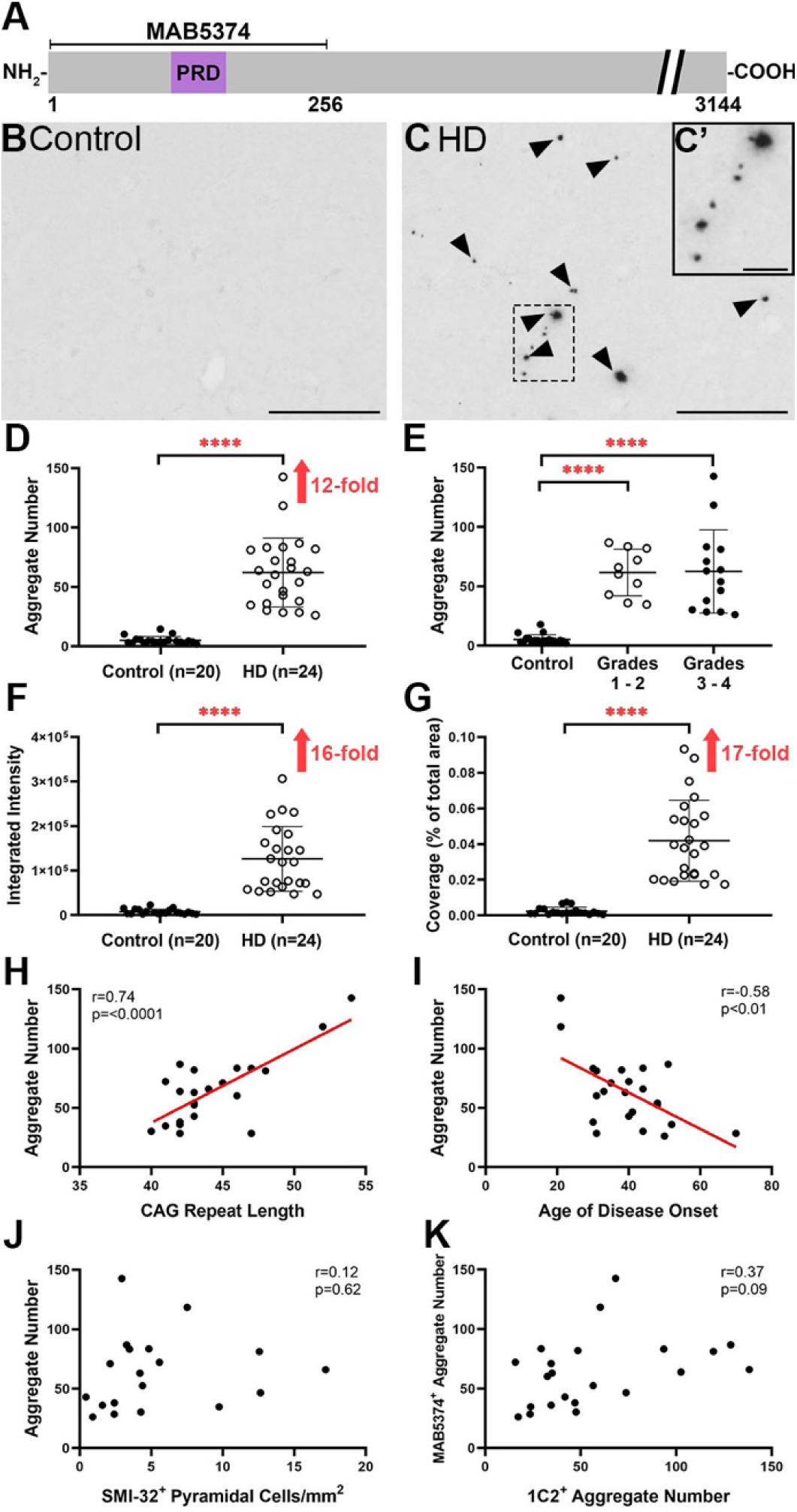
MAB5374 in cortical tissue microarray demonstrates Htt aggregate-specific immunoreactivity in HD cases. **(A)** MAB5374 binds to the N-terminal amino acid sequence 1-256 with the deletion of the polyQ tract of the Htt protein. **(B-C)** Representative images of MAB5374 immunoreactivity in (B) control and (C) HD MTG cores from the HD TMA. Black arrowheads denote MAB5374^+^ aggregates. **(D)** High-throughput analysis of the TMA revealed a 12-fold increase in the number of MAB5374^+^ aggregates (p<0.0001), in HD cores (n=24) relative to control cores (n=20). **(E)** Subdividing HD cases according to Vonsattel striatal neuropathological grade revealed a 11.4-fold increase in MAB5374^+^ aggregate number (p<0.0001) for Grades 1-2 HD cases (n=10) compared with controls (n=20) and a 11.5-fold increase (p<0.0001) in Grades 3-4 HD cases (n=14) relative to controls. However, there was no significant difference in aggregate number between Grades 1-2 and Grades 3-4 HD cases (p>0.9999). **(F-G)** A (F) 16-fold increase in MAB5374 integrated intensity (staining expression) (p<0.0001) and (G) 17-fold increase in MAB5374 coverage (p<0.0001) was observed in HD cores (n=24) relative to control cores (n=20). **(H)** A strong positive correlation was found between MAB5437^+^ aggregate number and CAG repeat length (n=22; r=0.74; p<0.0001). **(I)** A moderate negative correlation was found between MAB5437^+^ aggregate number and age of disease onset (n=23; r=-0.58; p=<0.01). **(J-K)** No significant correlation was found between MAB5374^+^ aggregate number and (J) SMI-32^+^ pyramidal cell density (n=19; r=0.12; p=0.62), and (K) 1C2^+^ aggregate number (n=22; r=0.37; p=0.09). Significance values are based on *p*-values from a two-tailed unpaired *t*-test for D, F, and G, and from a Kruskal-Wallis test with Dunn’s multiple comparisons test for E. \*\*\*\**p* < 0.0001. Scale bar corresponds to 100 μm in B and C, and 20 μm in C’. PolyQ = polyglutamine tract; PRD = proline-rich domain; HD = Huntington’s disease; Htt = Huntingtin; CAG = cytosine-adenine-guanine; SMI-32 = Sternberger monoclonal incorporated antibody 32.

To better understand the variation in data within the HD cohort, the number of MAB5374^+^ aggregates for each HD case were further subgrouped according to important HD clinico-pathological variables, including CAG repeat length and age of disease onset (Figure 5 H-K). MAB5374^+^ aggregates displayed a strong positive correlation with CAG repeat length (r=0.74; p<0.0001) and a moderate negative correlation with age of disease onset (r=-0.58; p=<0.01) (Figure 5 H-I). However, the abundance of MAB5374^+^ aggregates in HD was not related to SMI-32^+^ pyramidal cell loss or 1C2^+^ Htt deposition (Figure 5 J-K).

### MW1 demonstrates both diffuse and aggregated Htt immunoreactivity in HD cortical brain tissue

MW1 was generated using a GST-tagged recombinant protein containing an extended polyQ tract and 34 amino acids from the dentatorubralpalliodoluysian atrophy-causing gene (Figure 6 A) (Ko et al., 2001). MW1 immunoreactivity demonstrated punctate staining indicating Htt aggregates in HD tissue, in addition to diffuse Htt immunoreactivity in both HD and control cases (Figure 6 B-C). TMA analysis of MW1 immunoreactivity demonstrated a 2.5-fold increase in the number of MW1^+^ aggregates in HD relative to controls (p<0.001) (Figure 6 D). Subdividing the HD cases according to early (Grades 1-2) and advanced (Grades 3-4) neuropathological grade revealed no difference in MW1^+^ aggregates between HD Grades (Figure 6 E). Furthermore, there were no significant changes in MW1 integrated intensity and coverage in HD cores relative to controls (Figure 6 F-G). The number of MW1^+^ aggregates was moderately correlated with CAG repeat length (r=0.63; p<0.01), and with age of disease onset (r=-0.44; p<0.05) (Figure 6 H-I). However, the upregulation of MW1^+^ aggregates in HD was not related to SMI-32^+^ pyramidal cell loss or 1C2^+^ Htt deposition (Figure 6 J-K).

**Figure 6.**
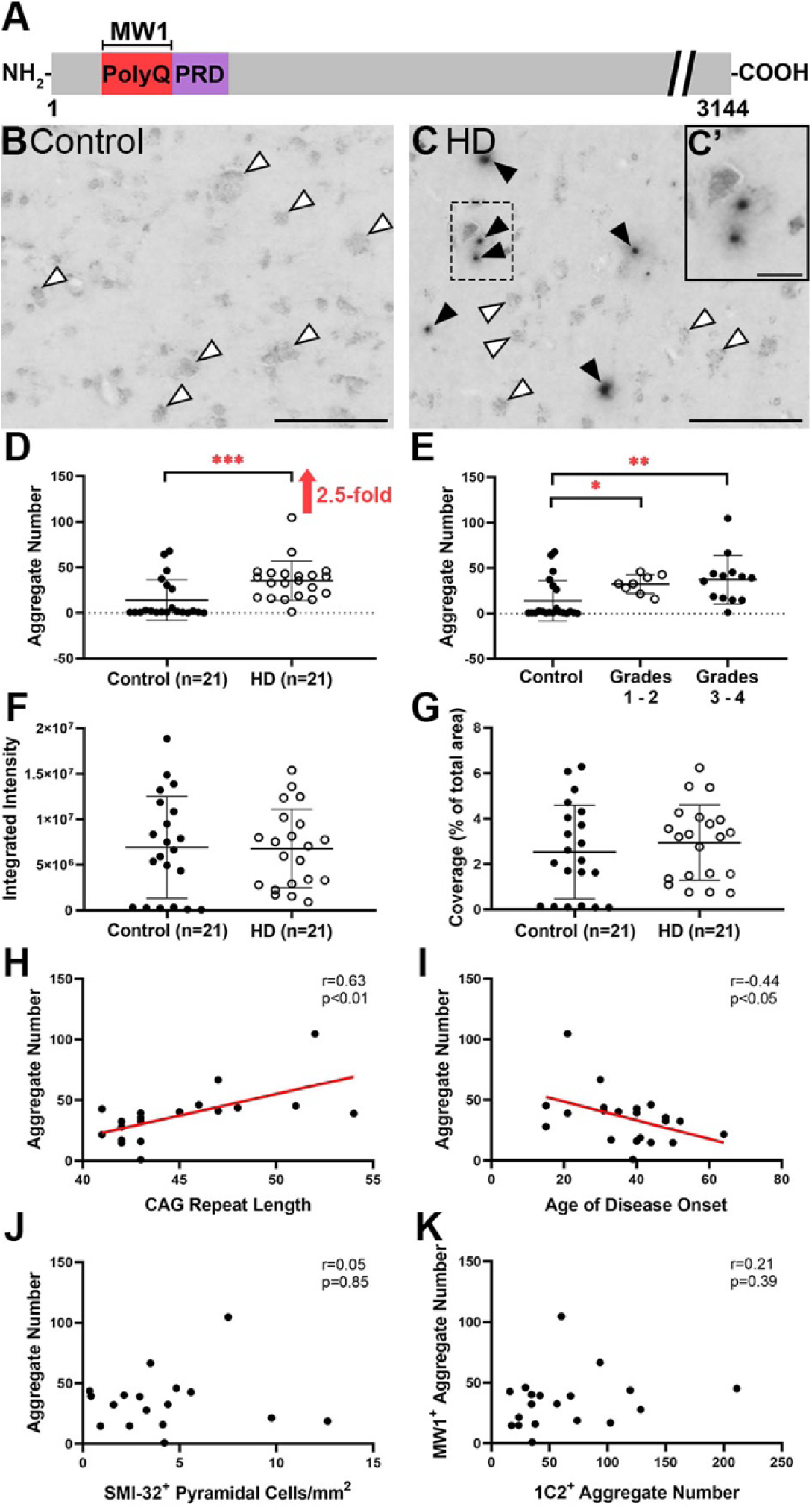
MW1 immunoreactivity in cortical tissue microarray demonstrates binding of both diffuse and aggregated Htt in HD cases. **(A)** Binding site of MW1 for the polyQ tract of Htt. **(B-C)** Representative images of MW1 immunoreactivity in (B) control and (C) HD cores from the TMA. Black arrowheads denote MW1^+^ aggregates in HD cases, as illustrated in the higher power inset (C’). White arrowheads denote diffuse MW1^+^ Htt in both control and HD cases. **(D)** TMA analysis revealed a 2.5-fold increase in the number of MW1^+^ aggregates (p<0.001) in HD cores (n=21) relative to control cores (n=21). **(E)** Subdividing HD cases according to neuropathological grade revealed a 2.3-fold increase in MW1^+^ aggregate number (p=0.05) between Grades 1-2 HD cases (n=8) relative to control (n=21), and a 2.7-fold increase (p=0.01) in Grades 3-4 HD cases (n=13) compared to controls. No significant difference between Grades 1-2 and Grades 3-4 HD cases. **(F-G)** There were no significant changes observed in (F) MW1 integrated intensity (p=0.84) and (G) MW1 coverage (p=0.52) in HD cores compared to control cores. (H) A moderate positive correlation was found between MW1^+^ aggregate number and CAG repeat length (n=19; r=0.63; p<0.01). **(I)** A moderate negative correlation was found between MW1^+^ aggregate number and age of disease onset (n=21; r=-0.44; p<0.05). **(J-K)** No significant correlation was found between MW1^+^ aggregate number and (J) SMI-32^+^ pyramidal cell density (n=17; r=0.05; p=0.85) and (K) 1C2^+^ aggregate number (n=19; r=0.21; p=0.39). Significance values are based on p-values from a two-tailed unpaired t-test for D, F, and G, and from a Kruskal-Wallis test with Dunn’s multiple comparisons test for E. \**p* < 0.05, \*\**p* < 0.01, \*\*\**p* < 0.001. Scale bar corresponds to 100 μm in B and C, and 20 μm in C’. PolyQ = polyglutamine tract; PRD = proline-rich domain; HD = Huntington’s disease; Htt = Huntingtin; CAG = cytosine-adenine-guanine; SMI-32 = Sternberger monoclonal incorporated antibody 32.

### EPR5526 demonstrates both diffuse and aggregated Htt immunoreactivity in HD cortical brain tissue with staining of pyramidal neurons

EPR5526 was generated against a synthetic peptide of the Htt N-terminal 100 amino acids and has been mapped to bind the Htt N17 domain, upstream of the polyQ tract (Figure 7 A) (Aviolat et al., 2019). EPR5526 immunoreactivity revealed staining patterns delineating pyramidal neurons in both control and HD tissue, in addition to punctate staining indicating Htt aggregates in HD cores (Figure 7 B-C). TMA analysis of EPR5526 immunoreactivity demonstrated a 4.9-fold increase in the number of EPR5526^+^ aggregates in HD cores (p<0.0001) (Figure 7 D). There was no difference in the number of EPR5526^+^ aggregates between HD Grades 1-2 and Grades 3-4 (Figure 7 E). Furthermore, there were no significant changes in EPR5526 integrated intensity or coverage in HD cores relative to controls (Figure 7 F-G). The number of EPR5526^+^ aggregates demonstrated a moderate positive correlation with CAG repeat length (r=0.47; p<0.05), and a moderate negative correlation with age of disease onset (r=-0.47; p=0.04) (Figure 7 H-I) but was not correlated with SMI-32^+^ pyramidal cell loss or 1C2^+^ Htt deposition (Figure 7 J-K).

**Figure 7.**
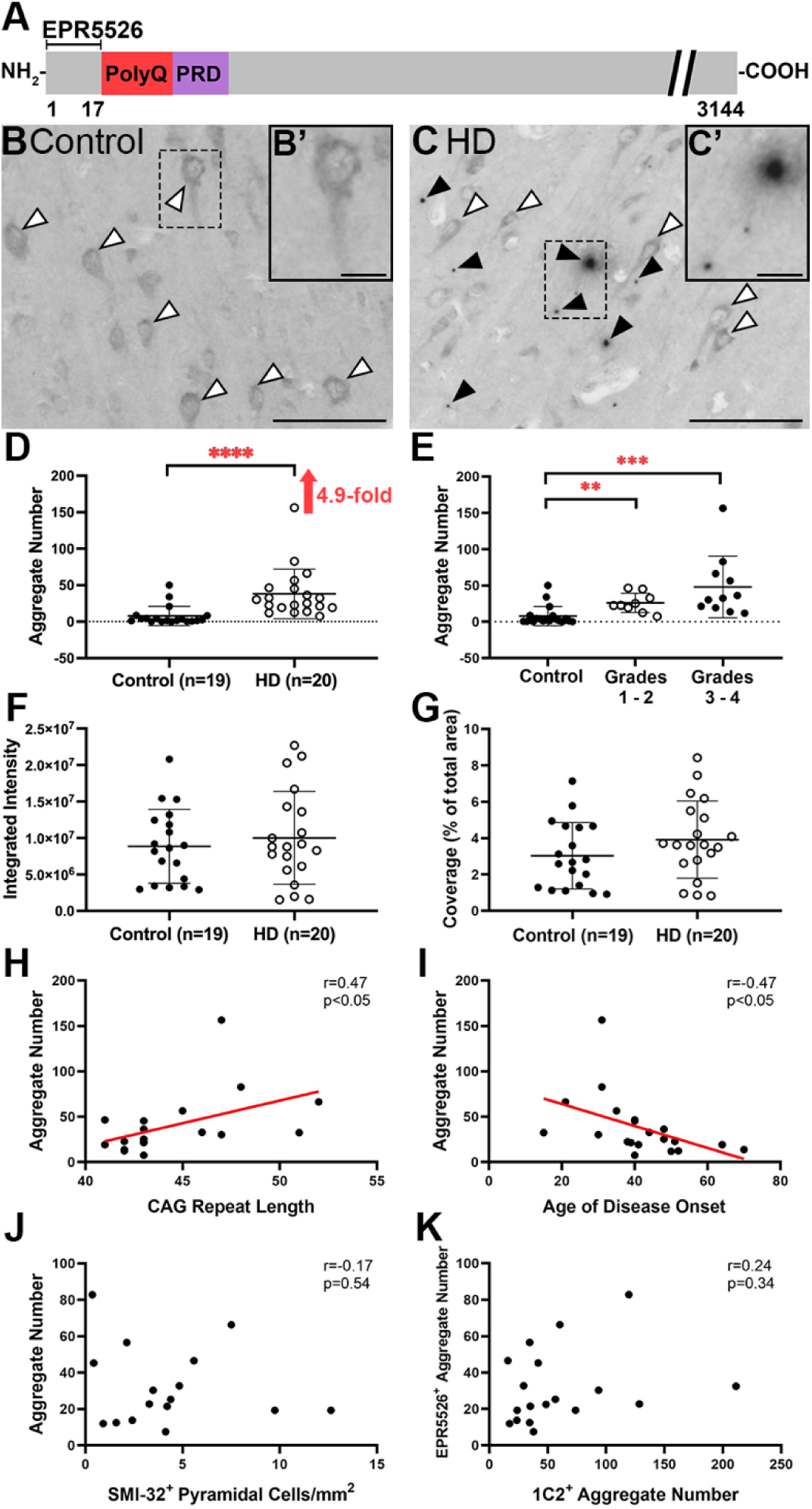
EPR5526 immunoreactivity in cortical tissue microarray demonstrates diffuse and aggregated Htt in HD cases. **(A)** Binding site of EPR5526 for the N-terminal amino acid sequence 1-100 of Htt. **(B-C)** Representative images of EPR5526 immunoreactivity in (B) control and (C) HD cores from the TMA. White arrowheads denote EPR5526^+^ diffuse Htt immunoreactivity outlining pyramidal neurons in both control and HD cases, as illustrated in the higher power inset (B’). Black arrowheads denote EPR5526^+^ aggregates in HD cases, as illustrated in (C’). **(D)** TMA analysis revealed a 4.9-fold increase in the number of EPR5526^+^ aggregates (p<0.0001) in HD cores (n=20) relative to control cores (n=19). **(E)** Subdividing HD cases according to neuropathological grade revealed a 3.3-fold increase in EPR5526^+^ aggregate number (p<0.01) for Grades 1-2 HD cases (n=9) compared with controls (n=19), and a 6.1-fold increase (p<0.001) in Grades 3-4 HD cases (n=11) versus controls. There was no significant difference between Grades 1-2 and Grades 3-4 HD cases. **(F-G)** There were no significant changes observed in (F) EPR5526 integrated intensity (p=0.73), and (G) EPR5526 coverage (p=0.20) in HD cores compared to control cores. **(H)** A moderate positive correlation was found between EPR5526^+^ aggregate number and CAG repeat length (n=18; r=0.47; p<0.05). **(I)** A moderate negative correlation was found between EPR5526^+^ aggregate number and age of disease onset (n=20; r=-0.47; p<0.05). **(J-K)** No significant correlation was found between EPR5526^+^ aggregate number and (J) SMI-32^+^ pyramidal cell density (n=16; r=-0.17; p=0.54), and (K) 1C2^+^ aggregate number (n=18; r=0.24; p=0.34). Significance values are based on p-values from a two-tailed unpaired t-test for D, F, and G, and from a Kruskal-Wallis test with Dunn’s multiple comparisons test for E. \*\**p* < 0.01, \*\*\**p* < 0.001, \*\*\*\**p* < 0.0001. Scale bar corresponds to 100 μm in B and C, and 20 μm in B’ and C’. PolyQ = polyglutamine tract; PRD = proline-rich domain; HD = Huntington’s disease; Htt = Huntingtin; CAG = cytosine-adenine-guanine; SMI-32 = Sternberger monoclonal incorporated antibody 32.

### MAB5492 demonstrates immunoreactivity patterns not specific to HD based on aggregate-like binding in both HD and control cortical brain tissue

MAB5492 was generated against a recombinant human Htt protein fragment containing the N-terminal 82 amino acids and has been mapped to bind the Htt proline-rich domain (PRD) (Figure 8 A) (Dehay et al., 2007). Interestingly, MAB5492 immunolabeled punctate objects in both control and HD tissue (Figure 8 C). MAB5492 also demonstrated the presence of sparse diffuse Htt immunoreactivity within both control and HD cohorts (Figure 8 B-C). TMA analysis of MAB5492 immunoreactivity revealed no significant changes in MAB5492^+^ aggregate number and immunolabel coverage in HD cores relative to controls (Figure 8 D, G). Furthermore, subdividing the HD cases according to neuropathological grade revealed no differences in MAB5492^+^ aggregate number with advancing disease grade (Figure 8 E). However, surprisingly, there was a significant 1.3-fold decrease in MAB5492 integrated intensity in HD compared to the highly variable control cohort (p=0.02) (Figure 8 F). It is evident that MAB5492 immunoreactivity within the control cohort is highly variable compared to the HD group, as evidenced by unequal variances (Figure 8 D-G). MAB5492^+^ aggregate number did not correlate with CAG repeat length, age of disease onset, SMI-32^+^ pyramidal cell loss or 1C2^+^ aggregates (Figure 8 H-K).

**Figure 8.**
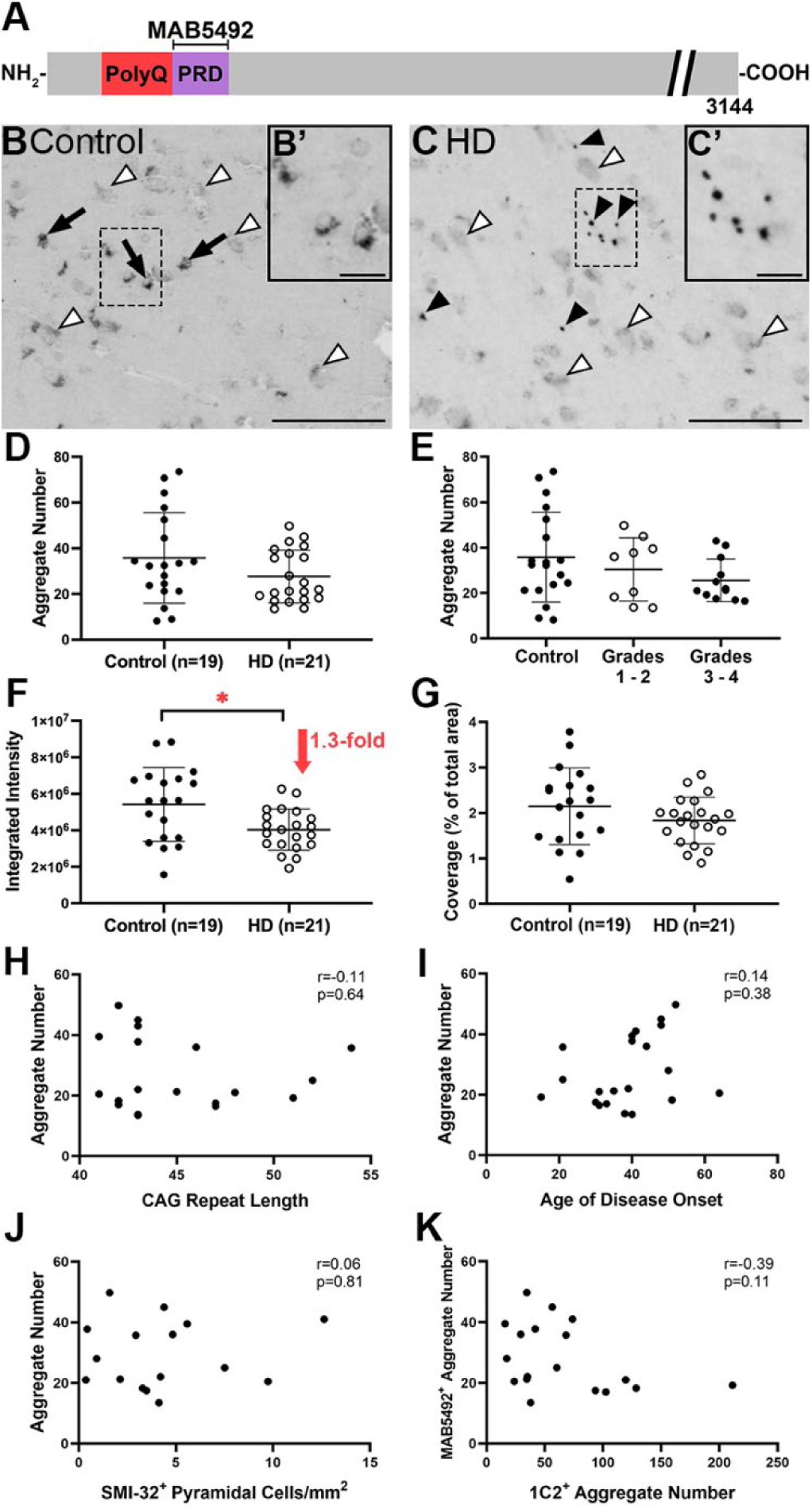
MAB5492 in cortical tissue microarray demonstrates immunoreactivity patterns not specific to HD based on aggregate-like binding in both HD and control human brain tissue. **(A)** Binding site of MAB5492 for the N-terminal amino acid sequence 1-82 of Htt. **(B-C)** Representative images of MAB5492 immunoreactivity in (B) control and (C) HD cores from the TMA. Black arrows denote MAB5492^+^ punctate Htt staining in control cases, as illustrated in the higher power inset (B’). Black arrowheads denote MAB5492^+^ aggregates in HD, as illustrated in (C’), and white arrowheads denote diffuse Htt in both control and HD cases. **(D)** TMA analysis revealed no significant changes were observed in the number of MAB5492^+^ aggregates (p=0.23) in HD cores (n=21) relative to control cores (n=19). **(E)** After subdividing HD cases according to neuropathological grade, there were no significant differences in MAB5492^+^ aggregate numbers between control cases (n=19), Grades 1-2 HD cases (n=9; p>0.99), and Grades 3-4 HD cases (n=12; p=0.50). **(F-G)** A significant 1.3-fold decrease in (F) MAB5492 integrated intensity was observed in HD cases compared to the control cohort (p=0.02), however no significant change was observed in MAB5492 coverage (p=0.21) in HD cores (n=21) relative to control cores (n=19). **(H-K)** No significant correlation was found between MAB5492^+^ aggregate number and (H) CAG repeat length (n=19; r=-0.11; p=0.64), (I) age of disease onset (n=21; r=0.14; p=0.38), (J) SMI-32^+^ pyramidal cell density (n=16; r=0.06; p=0.81), and (K) 1C2^+^ aggregate number (n=18; r=-0.39; p=0.11). Significance values are based on p-values from a two-tailed unpaired t-test. \**p* < 0.05. Scale bar corresponds to 100 μm in B and C, and 20 μm in B’ and C’. PolyQ = polyglutamine tract; PRD = proline-rich domain; HD = Huntington’s disease; Htt = Huntingtin; CAG = cytosine-adenine-guanine; SMI-32 = Sternberger monoclonal incorporated antibody 32.

### MW8 immunoreactivity demonstrates lack of specificity for aggregated Htt in HD cortical brain tissue

MW8 was generated against human Htt exon 1 containing a 67-residue polyQ tract, resulting in its specificity for Htt N-terminal amino acids 83-90 (Figure 9 A) (Ko et al., 2001). Qualitative examination of MW8 immunoreactivity demonstrated diffuse immunoreactivity localized to cell bodies in both HD and control tissue (Figure 9 B-C). However distinct aggregate staining was not observed and therefore the number of aggregates was not quantified (Figure 9 B-C). Analysis of MW8 immunoreactivity in the TMA revealed an increase in MW8 immunolabel expression in HD, as evidenced by a 1.7-fold increase in MW8 integrated intensity (p<0.01) and 1.1-fold increase in MW8 coverage (p<0.01) in HD cores relative to controls (Figure 9 D, F). There was no difference in MW8 integrated intensity between HD Grades 1-2 and Grades 3-4 (Figure 9 E), and MW8 integrated intensity did not correlate with any clinico-pathological variables (Figure 9 G-J).

**Figure 9.**
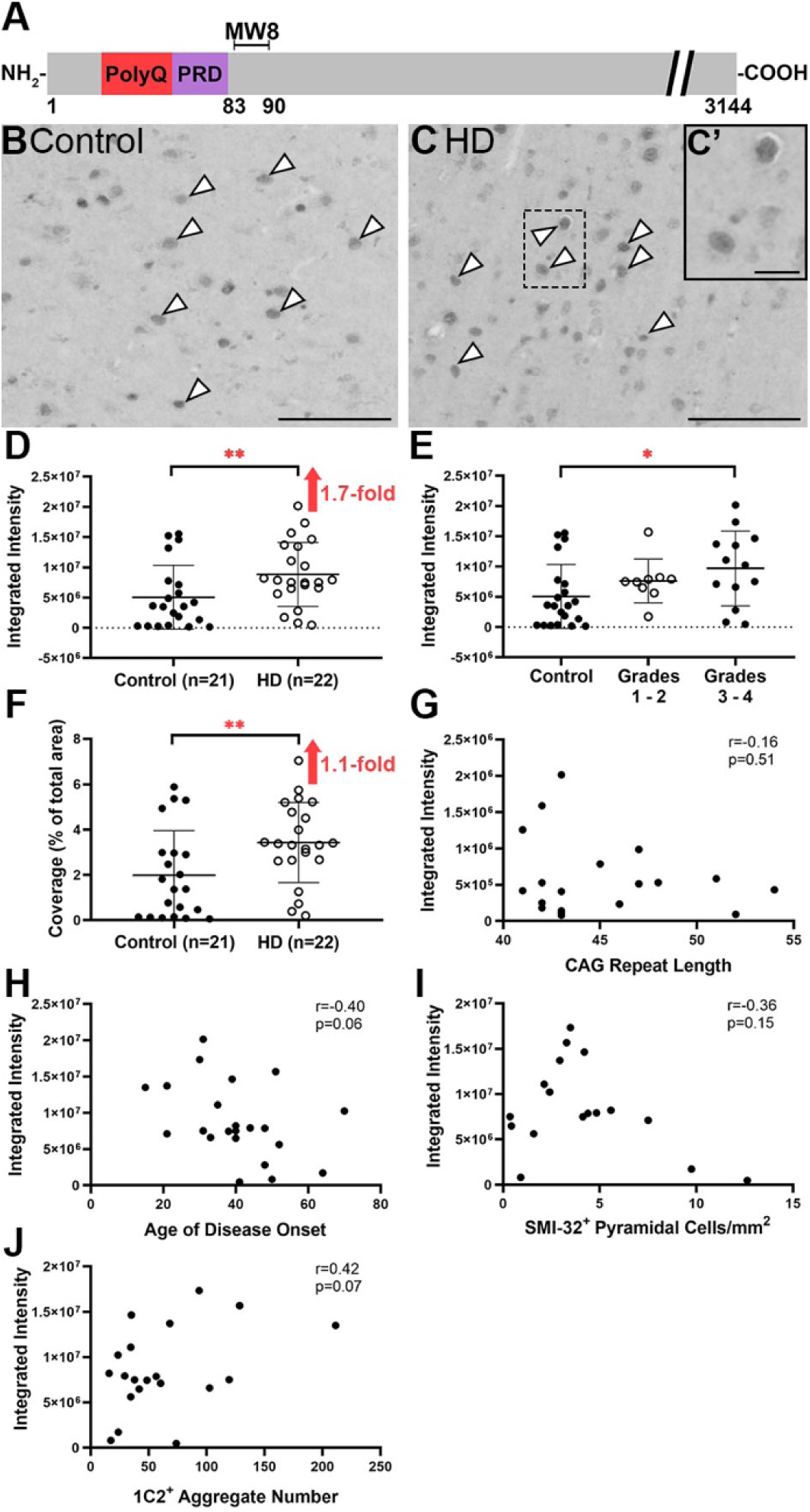
MW8 immunoreactivity in cortical tissue microarray demonstrates lack of specificity for aggregated Htt in HD. **(A)** Binding site of MW8 for the N-terminal amino acid sequence 83-90 of Htt. **(B-C)** Representative images of MW8 immunoreactivity in (B) control and (C) HD cores from the TMA. White arrowheads denote MW8^+^ Htt immunoreactivity localized to the cell body in both control and HD cases. **(D)** TMA analysis revealed a 1.7-fold increase in MW8 integrated intensity (staining expression) (p<0.01) in HD (n=22) relative to control cores (n=21). **(E)** After subdividing HD cases according to neuropathological grade, there were no significant differences in MW8 expression (p=0.76) between control cases (n=21) and Grades 1-2 HD cases (n=9). However there was a 1.9-fold increase in MW8 expression (p<0.05) in Grades 3-4 HD cases (n=13) compared to controls. There was no significant difference in MW8 staining expression for Grades 1-2 versus Grades 3-4 HD cases. **(F)** A 1.1-fold increase was observed in MW8 expression (p<0.01) in HD compared to control cores. **(G-J)** No significant correlation was found between MW8 expression and (G) CAG repeat length (n=20; r=-0.16; p=0.51), (H) age of disease onset (n=22; r=-0.40; p=0.06), (I) SMI-32^+^ pyramidal cell density (n=17; r=-0.36; p=0.15), and (J) 1C2^+^ aggregate number (n=20; r=0.42; p=0.07). Significance values are based on p-values from a two-tailed unpaired t-test. \**p* < 0.05, \*\**p* < 0.01. Scale bar corresponds to 100 μm in B and C, and 20 μm in C’. PolyQ = polyglutamine tract; PRD = proline-rich domain; HD = Huntington’s disease; Htt = Huntingtin; CAG = cytosine-adenine-guanine; SMI-32 = Sternberger monoclonal incorporated antibody 32.

### 2B7 demonstrates neuropil-like immunoreactivity in control and HD cortical brain tissue with a lack of specificity for aggregated Htt in HD

2B7 was generated against the human Htt protein to result in its specificity for Htt N-terminal amino acids 7-13 (Figure 10 A) (Weiss et al., 2009). 2B7 revealed interesting neuropil-like immunoreactivity in both HD and control tissue, however, distinct aggregate staining was not observed and therefore the number of aggregates were not quantified (Figure 10 B-C). TMA analysis of 2B7 immunoreactivity revealed no significant differences in 2B7 integrated intensity and coverage in HD cores relative to controls, and no association between 2B7 integrated intensity and neuropathological grade (Figure 10 D-F). 2B7 integrated intensity did not correlate with any clinico-pathological variables (Figure 10 G-J).

**Figure 10.**
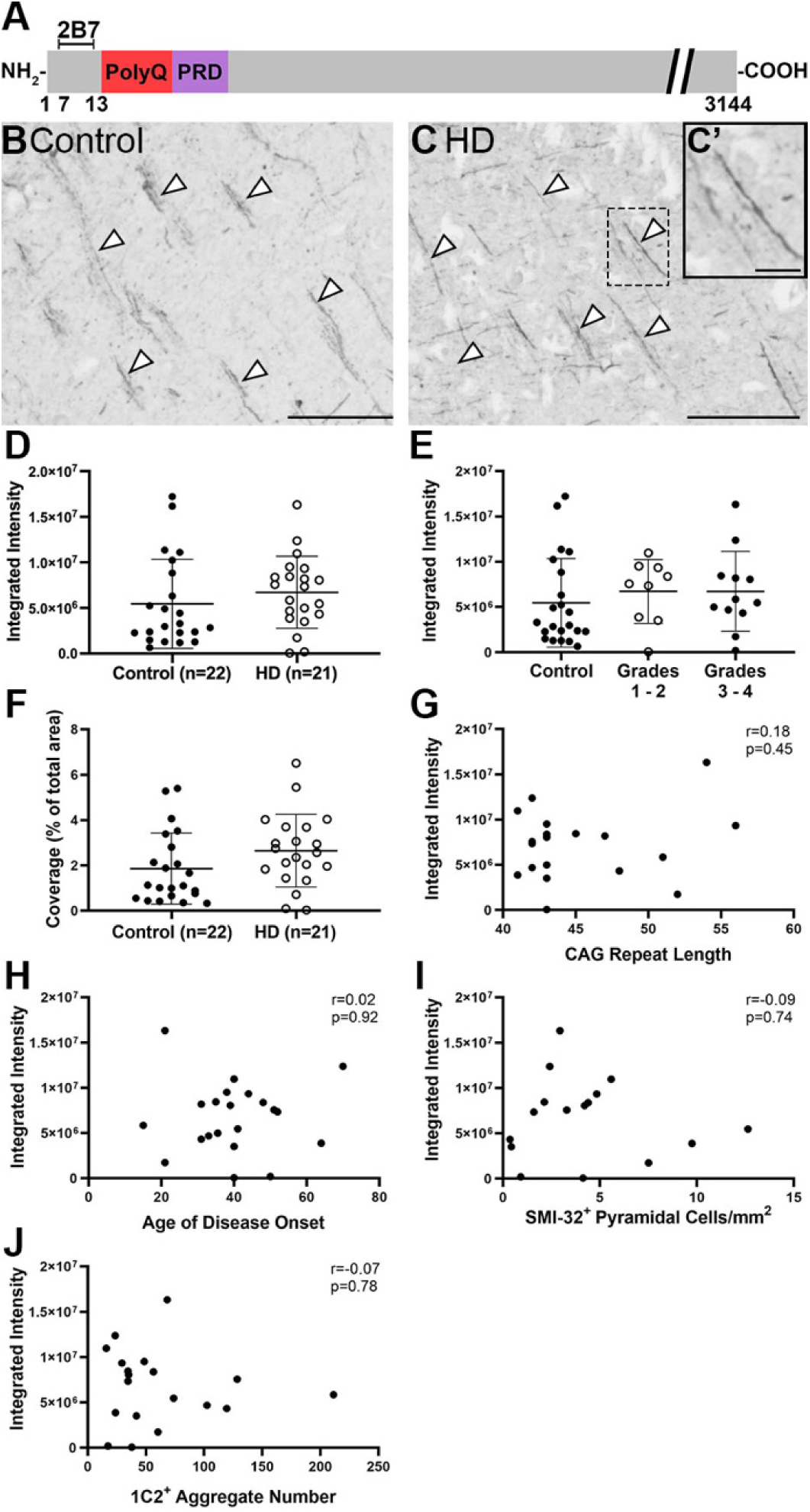
2B7 in cortical tissue microarray demonstrates neuropil-like immunoreactivity in control and HD human brain tissue with a lack of specificity for aggregated Htt. **(A)** Binding site of 2B7 for the N-terminal amino acid sequence 7-13 of Htt. **(B-C)** Representative images of 2B7 immunoreactivity in (B) control and (C) HD cores from the TMA. White arrowheads denote 2B7^+^, neuropil-like Htt staining in both control and HD cases. **(D-F)** TMA analysis revealed no significant changes in (D) 2B7 integrated intensity (staining expression) (p=0.18) and (F) 2B7 coverage (p=0.07) in HD cores (n=21) relative to control cores (n=22). **(E)** After subdividing HD cases according to neuropathological grade, there were no differences in 2B7 expression between control cases (n=22) versus Grades 1-2 HD cases (n=9; p=0.76), and Grades 3-4 HD cases (n=12; p=0.85). There was also no significant difference between 2B7 expression for Grades 1-2 versus Grades 3-4 HD cases. **(G-J)** No significant correlation was found between 2B7 expression and (G) CAG repeat length (n=19; r=0.18; p=0.45), (H) age of disease onset (n=21; r=0.02; p=0.92), (I) SMI-32^+^ pyramidal cell density (n=16; r=-0.09; p=0.74), and (J) 1C2^+^ aggregate number (n=19; r=-0.07; p=0.78). Significance values are based on *p*-values from a two-tailed unpaired *t*-test. Scale bar corresponds to 100 μm in B and C, and 20 μm in C’. PolyQ = polyglutamine tract; PRD = proline-rich domain; HD = Huntington’s disease; Htt = Huntingtin; CAG = cytosine-adenine-guanine; SMI-32 = Sternberger monoclonal incorporated antibody 32.

### Post-mortem delay and age at death do not impact TMA immunoreactivity data except for MAB5492

To determine the impact of post-mortem delay (PMD) and age at death on the data generated for each case, the relationship was examined between the integrated intensity (as a measure of staining expression) for each marker, and both PMD and age at death, respectively. Integrated intensity was utilized for PMD and age at death correlations as this variable was measured for all immunolabels investigated. To rule out the influence of disease, the integrated intensity data was pooled for both control and HD cases for these comparisons. The strength of the correlation was classified according to the criteria in Table 5, and correlations were considered significant if there was a *p*-value of ≤ 0.05. For all markers, with the exception of MAB5492, there was no significant correlation between integrated intensity and either PMD or age at death (Table 6). A moderate correlation was found between MAB5492 integrated intensity and age at death (r=0.52; p<0.001; Figure 11), thereby suggesting that aging may be a factor influencing MAB5492 immunoreactivity as opposed to CAG repeat length, age of onset, or PMD.

**Figure 11.**
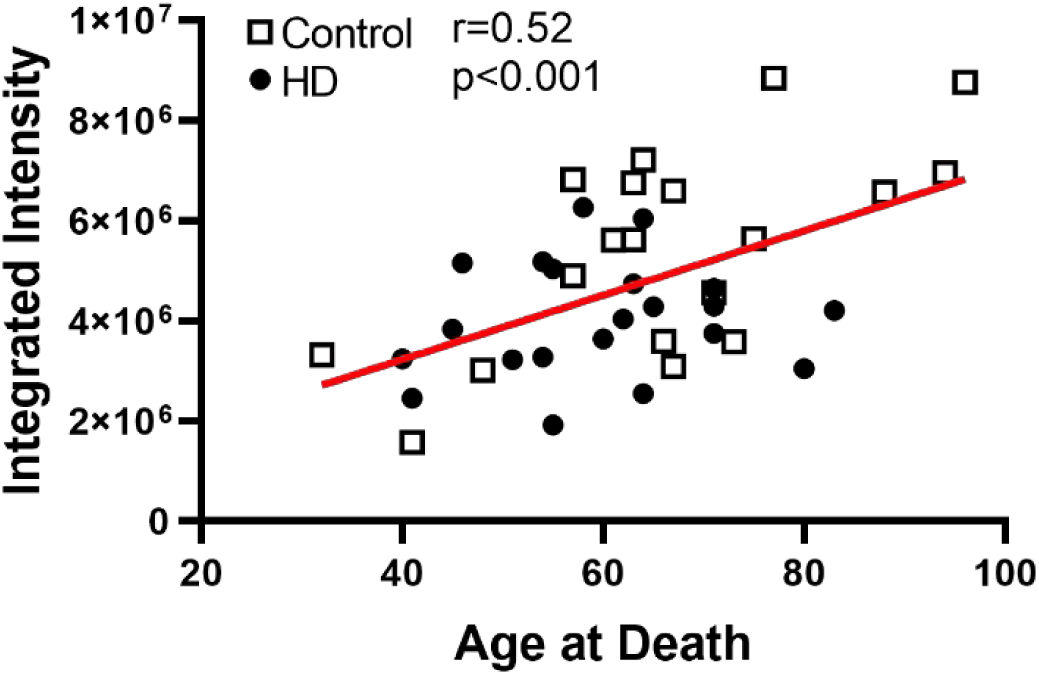
Correlation between MAB5492 integrated intensity and age at death. Within the pooled control and HD cohort (n=40), a moderate positive correlation was found between MAB5495 protein expression (integrated intensity) and age at death (r=0.52; p<0.001).

**Table 6.**
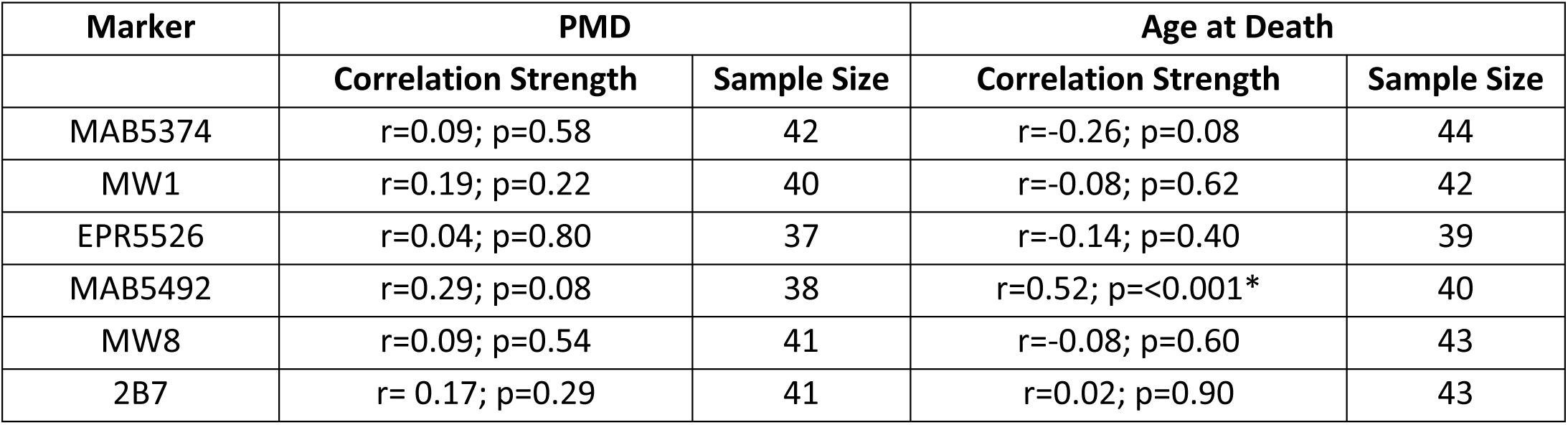
Correlations between marker expression with post-mortem delay and age at death. Details include the parameter used to correlate with post-mortem delay and age at death for each marker, the *r* and *p* values obtained using a Spearman’s test, and the sample size for each comparison comprising of pooled HD and control cases. * denotes statistically significant correlation. PMD = post-mortem delay.

| Marker | PMD |  | Age at Death |  |
| --- | --- | --- | --- | --- |
|  | Correlation Strength | Sample Size | Correlation Strength | Sample Size |
| MAB5374 | $r=0.09$; $p=0.58$ | 42 | $r=-0.26$; $p=0.08$ | 44 |
| MW1 | $r=0.19$; $p=0.22$ | 40 | $r=-0.08$; $p=0.62$ | 42 |
| EPR5526 | $r=0.04$; $p=0.80$ | 37 | $r=-0.14$; $p=0.40$ | 39 |
| MAB5492 | $r=0.29$; $p=0.08$ | 38 | $r=0.52$; $p<0.001^*$ | 40 |
| MW8 | $r=0.09$; $p=0.54$ | 41 | $r=-0.08$; $p=0.60$ | 43 |
| 2B7 | $r=0.17$; $p=0.29$ | 41 | $r=0.02$; $p=0.90$ | 43 |

## Discussion

This study is the first to screen a diverse range of commercially available and novel anti-Huntingtin (Htt) antibodies using a well-established human brain tissue microarray (TMA) platform (Singh-Bains et al., 2021). This study profiled Htt immunoreactivity in TMAs comprising the largest sample size of both Huntington’s disease (HD) and matched control human brains, allowing high-throughput analysis of Htt aggregates which were correlated with measures of HD clinicopathology. The results of this study support the initial hypothesis, that utilizing a combination of different anti-huntingtin (Htt) antibodies will increase our chances of capturing different types of Htt in the human HD brain. Each anti-Htt antibody resulted in a unique pattern of immunoreactivity, and some antibodies unexpectedly immunolabeled non-aggregate features in both control and HD cases. Htt aggregates were more readily detected using N-terminal anti-Htt antibodies compared to C-terminal antibodies, providing further evidence that N-terminal polyglutamine (polyQ)-containing fragments are the major component of Htt pathological aggregates. Most importantly, the results suggested that a greater presence of Htt aggregates is not associated with the degree of cortical pyramidal neuronal loss in HD.

### Markers for Htt aggregates: MAB5374, MW1, and EPR5526

In this study, three out of the twelve anti-Htt antibodies trialed with human brain IHC resulted in immunolabelling of Htt aggregates, including MAB5374, MW1, and EPR5526. Each aggregate-detecting antibody binds Htt within the N-terminal region of the protein, which is consistent with earlier immunohistochemical studies of the human HD brain that observe aggregates exclusively with N-terminal directed antibodies (Becher et al., 1998; DiFiglia et al., 1997; Sieradzan et al., 1999).

The marker with the greatest specificity for immunolabelling Htt aggregates was MAB5374, which binds the N-terminal 256 amino acids of Htt protein, and is the only antibody included in this study that has been extensively utilized for human brain IHC. Upon visual inspection of immunolabeled tissue, MAB5374^+^ aggregates were distinct, punctate objects in HD tissue, compared to minimal immunoreactivity observed in control tissue (Figure 5 B-C). This staining pattern was reflected in the quantitative analysis of cortical HD and control TMA cores, with a significant increase in MAB5374^+^ aggregate number, integrated intensity (staining expression), and immunolabel coverage (expressed as a percentage of total core area) observed in HD compared to control cases (Figure 5 D, F, G). These results align with previously published anatomical studies utilizing MAB5374 or the clone EM48, reporting a significant increase in mutant Htt aggregates in the insular cortex, parietal lobe, frontal lobe, and motor cortex of the HD brain (Gutekunst et al., 1999; Kuemmerle et al., 1999; Schilling et al., 2007; van Roon-Mom et al., 2006). Most of these studies observe EM48^+^ or MAB5374^+^ aggregates localized to the neuropil and cytoplasm of cortical neurons, however, Schilling et al. (2007) additionally observed EM48^+^ aggregates within the nucleus (Gutekunst et al., 1999; van Roon-Mom et al., 2006). With only one previous study identifying nuclear MAB5374^+^ aggregates, there is a need for further investigation into the subcellular localization of MAB5374 immunoreactivity to determine what fraction of the total (cytoplasmic + nuclear) Htt aggregates are immunolabeled. If MAB5374 is unable to consistently capture nuclear Htt aggregates in the human brain, the antibody may need to be combined with an additional nuclear-directed anti-Htt antibody for future studies profiling Htt deposition.

### Total Htt detection with MW1 and EPR5526

Unlike MAB5374, antibodies MW1 and EPR5526 immunolabeled both normal and aggregated Htt in the human brain cortical TMAs. Antibodies MW1 and EPR5526 target extreme N-terminal domains of the Htt sequence, whereas MAB5374 targets the longer N-terminal 1-256 amino acids thereby reinforcing the notion that N-terminal fragments of different lengths are contributing to the formation of Htt aggregates (Kolla et al., 2021). The heterogeneity of both the composition and structure of Htt protein aggregates throughout the HD brain points to the need for an expanded panel of antibodies targeting different regions of the protein to accurately reflect pathological Htt protein deposition in the HD brain.

In addition to immunolabelling Htt aggregates, MW1 displayed diffuse Htt immunoreactivity (Figure 6 B-C), while EPR5526 immunolabeled the borders of pyramidal neurons (Figure 7 B-C) in both control and HD tissue. Quantitative analysis of HD and control cores demonstrated a significant increase in the number of Htt aggregates in HD for both antibodies (Figure 6 D; Figure 7 D). However, the integrated intensity and coverage of MW1 and EPR5526 immunoreactivity (consisting of diffuse and punctate staining) were highly variable within the control and HD cohorts, and did not demonstrate a significant change in HD compared to control (Figure 6 F-G; Figure 7 F-G). These results suggest that the overall Htt expression levels are not altered in the HD brain, therefore Htt aggregation is not associated with protein overexpression, which may reinforce the theory of Htt aggregation being the byproduct of a gain of toxic function by mutant Htt in the HD brain (Aronin et al., 1995; Scherzinger et al., 1999; Schulte & Littleton, 2011).

To date, this is the first study to profile MW1 in human HD brain tissue using IHC. MW1 is a widely used antibody in rodent models of HD and for detecting Htt expression in humans via Meso Scale Discovery (MSD) or Time-resolved Förster resonance energy transfer (TR-FRET) analysis of human brain lysates (Caron et al., 2021; Cicchetti et al., 2014; Legleiter et al., 2009; Southwell et al., 2015). MSD and TR-FRET utilizing MW1 are currently used in clinical trials to test the effects of novel Htt-lowering treatments, however, MW1 may not provide an accurate representation of Htt expression levels (Fodale et al., 2017). Multiple studies have noted the specificity of MW1, binding the Htt polyQ tract, as a limitation for objective Htt quantification due to the heterogeneity of this region (Aviolat et al., 2019; Fodale et al., 2017). A polyQ length-dependent bias was demonstrated to affect Htt detection levels via MSD, with a longer polyQ resulting in a higher rate of detection, and even small variations in polyQ length produced large inaccuracies in protein expression quantification (Aviolat et al., 2019). Considering the instability of the CAG repeat region and its propensity to undergo somatic repeat expansions in the human brain, polyQ-independent methods of Htt quantification are essential to gain an accurate representation of the protein in the human HD brain (Cariulo et al., 2017; Fodale et al., 2017). Furthermore, MW1 was raised against a construct containing an extended polyQ tract and 34 amino acids of the DRPLA-causing gene, another CAG trinucleotide repeat expansion disease. Therefore, MW1 is not strictly specific for human Htt protein and may immunolabel other polyQ-containing proteins associated with Htt aggregates, leading to a misrepresentation of Htt distribution by using this antibody.

Meanwhile, EPR5526 has been mapped to bind the N17 domain of Htt upstream of the polyQ tract, and may serve as a more aggregate-specific alternative to MW1 (Aviolat et al., 2019). The EPR5526^+^ immunoreactivity delineating pyramidal cells provides a preliminary indication of the subcellular localization of EPR5526^+^ aggregates within pyramidal neurons, while a subset of EPR5526^+^ aggregates appeared to be located extracellularly (Figure 7 C). At this stage, we cannot conclude that all pyramidal cells are immunolabelled with EPR5526, raising the need for further colocalization studies using a range of antibodies for both neuronal and non-neuronal cell markers to investigate the cellular distribution of EPR5526 within both pyramidal cells and other cell types. In the current literature, Htt aggregates are typically reported in the nucleus, cytoplasm, or the processes of neurons in the post-mortem human HD brain (DiFiglia et al., 1997; Gutekunst et al., 1999; Jansen et al., 2017; Sieradzan et al., 1999). However, extracellular protein aggregates have been observed in other neurodegenerative diseases, for example tau and alpha synuclein in Alzheimer’s disease and Parkinson’s disease respectively, suggesting that this phenomenon could also be occurring in the HD human brain (Emmanouilidou et al., 2011; Yamada, 2017). Moreover, mutant Htt aggregates were identified within the extracellular matrix of striatal allografts implanted in the brains of three HD patients, suggesting that Htt aggregates spread from the host brain to the transplanted tissue (Cicchetti et al., 2014). Therefore, true extracellular Htt aggregates may provide evidence of the transmission of Htt aggregates throughout the brain in HD

### Aggregate number correlates with CAG repeat length and age of disease onset but not striatal neuropathological grade

The relationships between the number of Htt aggregates and key clinicopathological variables associated with HD were investigated to provide insight into the contribution of Htt to HD pathology. In agreement with previously published studies of the human HD brain, Htt aggregate numbers determined using MAB5374, MW1, and EPR5526 did not correlate with striatal neuropathological grade, indicating that Htt aggregates within the cerebral cortex are not associated with the degree of striatal neurodegeneration in HD (Figure 5 E; Figure 6 E; Figure 7 E) (Becher et al., 1998).

However, this study demonstrated a significant positive correlation between Htt aggregate numbers detected with antibodies MAB5374, MW1, and EPR5526 and CAG repeat length (Figure 5 H; Figure 6 H; Figure 7 H). This relationship was first observed in an immunohistochemical study by Becher et al. (1998), utilizing post-mortem human brain tissue from 20 HD patients, and subsequent studies support the conclusion that Htt deposition does not correlate with Vonsattel striatal neuropathological grade and correlates more closely with CAG repeat length (Gutekunst et al., 1999; Kuemmerle et al., 1999). This relationship can be explained using evidence obtained from cell culture and animal models of HD on the dynamics of Htt aggregation, which demonstrate that Htt aggregation is dependent on CAG repeat length *in vitro* and *in vivo*, with a longer a polyQ tract resulting in a higher propensity to form strong hydrogen bonds and accumulate in aggregates (Legleiter et al., 2010; Perutz, 1996; Scherzinger et al., 1999; Vieweg et al., 2021). However, due to the lack of association between neuronal cell loss and Htt aggregates in this current study, it is still unclear whether a longer CAG repeat length, therefore a longer polyQ tract, confers a greater toxicity to the cell in the HD human brain.

Comparisons between the number of Htt aggregates detected using antibodies MAB5374, MW1, and EPR5526 versus the age of disease onset revealed, for the first time in human brain tissue, a significant inverse relationship in this study. Although there are no studies published investigating this relationship in the human HD brain, a correlation was recently demonstrated between Htt aggregate formation and symptom onset in a mouse model of HD (Landles et al., 2020). The potential applications of this finding were further highlighted when a negative correlation was found between mutant Htt in the cerebrospinal fluid of late-stage HD patients and the age of disease onset, therefore Htt detection may be used as a diagnostic tool to predict the clinical onset of HD (Southwell et al., 2015). As previously reported and confirmed within our HD cohort (Supplementary Figure 1) there is a well-established negative relationship between CAG repeat length and the age of disease onset, and the results from this study suggest a critical role for Htt aggregates in the underlying pathological mechanism driving HD clinical presentation (Andrew et al., 1993; Duyao et al., 1993; Halliday et al., 1998).

### MAB5374, MW1, and EPR5526 aggregate numbers do not correlate with pyramidal cell loss or 1C2^+^ aggregates

There was no significant relationship between the number of MAB5374^+^, MW1^+^ or EPR5526^+^ Htt aggregates and the density of SMI-32^+^ pyramidal cells in the MTG TMA, which suggests the number of aggregates does not correlate with pyramidal cell loss (Figure 5 J; Figure 6 J; Figure 7 J). This is the first study to report this relationship in the temporal cortex and is consistent with previous studies of other brain regions in the human HD brain, reporting highly variable aggregate numbers in the insular cortex irrespective of pyramidal cell loss and striatal neuropathological grade (Gutekunst et al., 1999; Kuemmerle et al., 1999). The MTG is one of the least degenerated regions of the cortex in HD, with stereological studies from our laboratory reporting a 33% loss of pyramidal cells in HD (Nana et al., 2014). Therefore, investigating this region may provide insight into the earlier events of HD pathogenesis proceeding extensive neuronal loss, starting with the accumulation of Htt (Nana et al., 2014).

To further investigate the role of Htt in HD, an expanded toolset to accurately detect Htt is essential. 1C2 is a commonly used antibody in the HD literature for staining Htt aggregates, however, there were no significant correlations between the number of 1C2^+^ aggregates and Htt aggregates detected with MAB5374, MW1, and EPR5526 in the TMAs. This was a surprising result, particularly because MW1 has the same epitope as 1C2, binding the Htt elongated polyQ tract. However, 1C2 was raised against TATA-binding protein and is not strictly specific for human Htt protein. 1C2 immunoreactivity was profiled in the post-mortem HD human brain, where the antibody’s potential as a novel post-mortem diagnostic tool for HD was demonstrated due to the selective detection of aggregates in HD tissue (Herndon et al., 2009). However, the results from the present study lead to questioning the specificity of 1C2, considering the respective immunogens of MAB5374, MW1, and EPR5526 are more closely related to human Htt.

### MAB5492 is not suitable for accurate detection of human Htt

Despite the reported specificity of MAB5492 for the Htt N-terminus, the antibody failed to specifically detect Htt aggregates in the TMAs. Qualitative examination of MAB5492 immunoreactivity revealed punctate staining in both control and HD tissue (Figure 8 B-C), and there was no significant difference in the number of punctate objects between the two cohorts (Figure 8 D). However, punctate staining in the HD cores had a rounder, aggregate-like shape, whereas punctate staining in the control cores had a more irregular or crescent-like shape (Figure 8 B’, C’). The MAB5492 clone, 2B4, has been previously published in very few studies utilizing human HD brain tissue. The presence of cytoplasmic Htt has been reported in both control and HD human tissue using 2B4, with additional staining of nuclear and aggregated Htt in HD (Lunkes et al., 2002). However, 2B4 immunoreactivity was compared with 1C2 in 19 different human HD brains, and punctate Htt staining was also observed in control tissue using 2B4, leading to the conclusion that the antibody was unsuitable for post-mortem diagnosis of HD (Herndon et al., 2009). In rodent and cell culture models of HD, MAB5492 has been reported to be a reliable marker to detect mutant Htt aggregates using IHC, western blots, and immunoassays (Schindler et al., 2021; Schut et al., 2015). However, this study demonstrated the failure of MAB5492 to detect Htt aggregates in human HD brain tissue, suggesting that conclusions drawn with the use of this antibody for the detection of Htt in both animal and *in vitro* models of HD may not be translatable to the human brain. This could suggest that major differences exist between the biochemical and structural properties of Htt aggregates formed in the human brain compared to those observed in transgenic models or *in vitro*. For example, recent cryogenic electron microscopy studies from human brains displaying tauopathies and α-synucleinopathies have shown that the fibrils isolated from the brains of patients have vastly different core structures that those prepared from recombinant proteins (Arakhamia et al., 2020; Guerrero-Ferreira et al., 2018). Although the structure of Htt fibrils has not yet been determined, the differences may be attributed to the artificial constructs used for models containing a very long polyQ expansion (>90-1240 residues), which is beyond the physiological range for humans (Bayram-Weston et al., 2016). As a result, we need to determine a reliable toolset of anti-Htt antibodies which are specific to the human pathology to further elucidate the human disease state.

### MW8 immunolabeled soluble Htt in the human brain

Out of the six successful anti-Htt antibodies found to immunolabel human tissue in this study, MW8 displayed the greatest specificity for diffuse Htt. Qualitative analysis of MW8 immunoreactivity revealed diffuse, cytoplasmic staining without the presence of aggregates in both control and HD human tissue (Figure 9 B-C). Furthermore, a significant increase in MW8 integrated intensity was observed in HD cores compared to control cores, driven by higher staining expression in the Grades 3-4 HD cases (Figure 9 E-F). The absence of punctate staining was a surprising result because MW8 is considered a highly specific marker for Htt aggregates in rodent models of HD (Bayram-Weston et al., 2016; Ko et al., 2001; Landles et al., 2020). For example, MW8 was the superior Htt aggregate marker in a comparative study with four other commonly used anti-Htt antibodies including 1C2, MW1, and S830, across five different mouse models of HD (Bayram-Weston et al., 2016). Furthermore, MW8 has been utilized for immunoassays of human brain lysates to specifically detect aggregated Htt (Morozova et al., 2015; Reindl et al., 2019). The differences in IHC staining patterns may reflect the different conformations of Htt aggregates in the mouse and human brain respectively, with the MW8 epitope buried within aggregates in human tissue and inhibiting antibody binding (Khoshnan et al., 2013). Formic acid was utilized for antigen retrieval in this study to limit the effects of antigen masking, but the epitope of MW8 is very short (amino acids 83-90) and might have still been inaccessible within the structure of aggregated Htt. Meanwhile, MW8 successfully immunolabelled soluble Htt, and for the first time our results demonstrate that MW8 is not a suitable marker for Htt aggregates in human brain tissue. Further studies need to determine if different tissue preparation methods may influence the binding properties of this antibody, commencing with comparing paraffin embedded tissue IHC with fixed free-floating tissue IHC methods (Waldvogel et al., 2006).

### 2B7 is not suitable for detecting Htt aggregates in human HD brain tissue

The IHC staining pattern of 2B7 was unlike any of the anti-Htt antibodies included in this study, and different compared to other commercially available antibodies binding to a similar region of the Htt protein (DiFiglia et al., 1997; Sieradzan et al., 1999). Visual inspection of 2B7 immunoreactivity in the TMA revealed neuropil-like staining (Figure 10 B-C), and quantitative analysis revealed no difference in staining expression between the control and HD cohorts (Figure 10 D). These results were unexpected considering that 2B7 binds Htt within the N-terminal 17 amino acid (N17) domain and extensive evidence in the HD literature demonstrates the involvement of the Htt N-terminus in Htt aggregates (Becher et al., 1998; Cooper et al., 1998). Furthermore, previous studies have utilized antibodies directed to the Htt N17 domain to immunolabel nuclear Htt aggregates in HD human brain tissue and rodent HD models (Schilling et al., 2007). This is the first study to utilize 2B7 for IHC in human brain tissue, resulting in the novel observation that this antibody does not display diffuse or punctate immunoreactivity characteristic of normal and aggregated Htt, respectively. As mentioned above, 2B7 is commonly paired with MW1 for detecting Htt expression via immunoassays, with 2B7 used to capture total Htt expression (mutant + normal) (Fodale et al., 2017; Wild et al., 2015). However, considering the staining pattern observed in this study, combined with no significant difference in 2B7 integrated intensity between control versus HD cores, and no association with HD clinico-pathological variables, the specificity of 2B7 for human Htt is called into question. One possible explanation is that the Htt N17 domain could adopt a different conformation, or may be cleaved, or potentially undergo other post-translational modifications in the human brain which do not occur in preclinical models, preventing 2B7 from binding Htt. Different tissue preparation methods may also influence the binding properties of this antibody, and future studies should compare the binding patterns of 2B7 in paraffin embedded and fixed free-floating human brain tissue (Waldvogel et al., 2006). The results from this study demonstrate that 2B7 is unsuitable for detecting Htt aggregates in human HD brain tissue and also displays a potentially novel insight into the distribution of Htt within cellular processes in the MTG.

### C-terminal anti-Htt antibodies did not immunolabel Htt in human brain tissue

Of the twelve anti-Htt antibodies trialed in this study, six did not immunolabel human brain tissue. The epitopes of these six antibodies were all downstream of the polyQ tract, and included the three antibodies MAB2166, D7F7, and MAB2168 which label the C-terminus of Htt. This result can be justified for the antibody D7F7, which has previously been shown to react with human Htt expressed in mouse models of HD but not in human tissue, due to the differential processing of human Htt in the mouse versus the human brain (Franich et al., 2018). Previous studies suggest that antigen masking could have prevented the remaining antibodies from binding Htt at their respective epitopes, with that region of the protein buried inside an aggregate (Khoshnan et al., 2013). However, formic acid treatment was carried out for antigen unmasking to overcome this limitation, and most of these antibodies’ epitopes are downstream of the Htt N-terminus and thought to be excluded from aggregates (Schilling et al., 2007). The results from this study indicate the need to develop more effective antibodies directed to the Htt C-terminus to gain a true representation of the entire Htt protein in the human HD brain.

### Methodological Considerations

The TMAs utilized in this study served as a high-throughput screening platform to study Htt immunoreactivity in the post-mortem human brain. The TMAs contained a much larger number of human brain cases compared to the majority of HD human brain pathology studies to date, with a sample size of cortical cores containing up to 27 control and 28 HD cases. The TMA methodology conserves human tissue by only using a 7μm thick, 2mm diameter tissue samples per case on each TMA slide (Singh-Bains et al., 2021). Furthermore, one of the great advantages of TMA is that all cases immunolabeled for a particular marker of interest are included on a single glass slide, resulting in a high level of experimental standardisation for IHC conditions and imaging procedures.

However, the TMAs do provide some limitations to profile Htt immunoreactivity. Although the samples included in the TMA include cortical layers II-V, they may not represent the diversity of Htt staining in a larger tissue area. Furthermore, the 7μm thick sections result in limited anatomical detail, and future studies to investigate the subcellular localisation of Htt immunoreactivity should be conducted in thicker whole tissue sections to capture more complex aspects of cellular morphology.

Post-mortem human brain tissue was used in this study, with the advantage of serving as a relevant platform to study the human disease state. However, the main limitation of post-mortem tissue is that it is end-stage pathology. This limitation makes it difficult to determine the timeline of pathogenic events and draw causative relationships. For example, it is not possible to determine if Htt aggregates form early and promote neuronal survival or form late and induce cell death. The only way to address these questions in post-mortem tissue is to acquire tissue bequests from a significant sample of HD patients at time points prior to end stages (i.e. pre-symptomatic, or premature early causes of death) which is difficult. Attempting to address this limitation is extremely challenging, therefore studying tissue from the MTG, one of the less degenerated cortical regions of the HD brain, presumably reflecting earlier stages of pathology before the onset of severe cell loss is one of the approaches we use to investigate earlier disease indicators. Future studies should profile Htt immunoreactivity throughout multiple regions of the brain, thereby allowing the Htt aggregate data to be compared with the degree of heterogenous neuronal cell loss patterns to help explore relationships between Htt aggregates and neurodegeneration in more detail.

### Future Directions

The results from this preliminary screening study of Htt immunoreactivity in post-mortem human brain tissue have identified MAB5374, MW1, and EPR5526 as candidate antibodies to further investigate Htt aggregate localization in the human brain. Future studies will be conducted to determine the subcellular localization of Htt aggregates by carrying out multiplex fluorescence IHC in post-mortem human brain tissue and colabelling candidate aggregate markers with a range of cell types. These studies will further elucidate the differential localization of Htt aggregates within particular cellular compartments such as the nucleus versus the cytoplasm, to determine any associations between distribution location and important variables such as the degree of cortical pyramidal neuron loss. These future studies will reveal the presence or absence of extracellular Htt aggregates in HD, as well as the distribution of Htt aggregates within non-neuronal cell types.

Further biochemical analysis of the Htt species present in the human brain is also required to define the different forms of Htt that contribute to HD pathology, providing valuable information not only to explain the results of anatomical studies, but also to guide the design of future Htt-detecting tools.

## Conclusion

In conclusion, this study is the first to utilize human brain TMAs to screen a large range of anti-Htt antibodies in both HD and control human brain tissue. Antibodies MAB5374, MW1, and EPR5526 were identified to detect Htt aggregates in the human brain, with quantitative analysis revealing correlations between Htt aggregate deposition and both CAG repeat length and age of disease onset respectively, but not striatal neuropathological grade. Antibodies MAB5492, MW8, and 2B7 which are used frequently in animal and *in vitro* models of HD did not immunolabel human brain tissue, highlighting the differences in the properties of Htt aggregates that form in the human brain compared to those in preclinical models, and the need to develop human Htt-specific tools. Finally, 2E10, 4C9, D7F7, MAB5490, MAB2166, and MAB2168 trialed in this study did not display immunoreactivity in human brain tissue, prompting the need to develop better antibodies directed to the Htt C-terminus to investigate the contribution of the C-terminus to HD pathogenesis. This study has set up a platform to further investigate the distribution of Htt with respect to various cell types, other pathogenic proteins, and relevant interacting proteins throughout the human HD brain. Together, these future studies will aim to further elucidate the contribution of the Htt protein to HD pathology, improving our understanding of Htt as a key drug target for HD treatments.

## Ethics

All research procedures and protocols utilizing human brain tissue were conducted under full ethics approval from the New Zealand Health and Disability Ethics Committee (Ref.14/NTA/208) and with the informed consent from the families of the donors.

## Authors’ contributions

F.E.L. conducted the immunohistochemical staining, image acquisition, analysis of TMAs, figure production, draft manuscript production. A.Y.S.T. co-supervised F. E. L and contributed to immunohistochemical staining, TMA preparation, construction, imaging, analysis and construction of figures. N.F.M. contributed to the preparation and construction of TMAs used in this project. M.A.C. and R.L.M.F supervised the ethical collection, donor interaction, clinical assessment and processing of human brain tissue for the TMA platform. L.J.T conducted the retrospective examination of clinical symptomatology including determination of clinical age of disease onset. H.A.L. N.R. and L. A. and preformed antibody screening studies to select and prioritize the novel and commercial anti-huntingtin antibodies utilised for this project and which they generously provided. M.D. co-supervised F.E.L. and conceived, established and directs the TMA platform and high-content-analysis platform used in this study. M.K.S.B facilitated the project design, project implementation, direction, construction of TMAs, primary supervision of F.E.L and manuscript preparation. M.K.S.B, H.A.L, and R.L.M.F conceived the study concept. F.E.L. wrote the draft and completed the final manuscript with contributions edits and approvals of the final manuscript from N.F.M., M.A.C., L.J.T., N.R., L. A., R.L.M.F, M.D., H.A.L., and M.K.S.B.

## Acknowledgements

We express our appreciation to all donor families in New Zealand, who through their generosity have provided their invaluable tissue donation to the Neurological Foundation Human Brain Bank for the construction of HBTMAs. We thank Senior Histologist S. Amirapu for her contribution to tissue processing and paraffin embedding protocols for TMA preparation. We thank C. Turner for his expert neuropathological assessments of the cases utilized for TMA construction. We acknowledge the excellent work and assistance of M. Eszes (Human Brain Bank), K. Hubbard, R. Parker and P. Anscombe (Research Technicians). We acknowledge the staff at the Biomedical Imaging Research Unit (R. Kurian and P. V. Anekal) for VSlide Scanner support.

## Figure Legends

**Additional file 1: Supplementary Table 1.** Summary of successful and unsuccessful anti-Htt antibodies applied to paraffin-embedded HD and control human brain tissue using immunohistochemistry. Table includes all IHC conditions tested for each antibody: antibody concentration (up to four different concentrations were tested), antigen retrieval buffer, and DAB incubation time. All antibodies were treated with formic acid in addition to the antigen retrieval buffer listed. The optimal IHC conditions for the successful antibodies (highlighted in green, above dashed line) which displayed immunoreactivity are also included. The antibodies denoted with an asterisk immunolabeled Htt aggregates. IHC = immunohistochemistry; DAB = 3,3’-diaminobenzidine; TMA = tissue microarray; HD = Huntington’s disease; Htt = Huntingtin.

**Additional file 2: Supplementary Figure 1.** Correlation between CAG repeat length and age of disease onset within the HD TMA cohort (n=28). A strong negative correlation was found between the CAG repeat length and age of disease onset (r=-0.73; p=<0.0001).

## Abbreviations

CAG: Cytosine-adenine-guanine
DAB: Diaminobenzidine
H_2_O_2_: Hydrogen peroxide
HD: Huntington’s disease
Htt: Huntingtin
IHC: Immunohistochemistry
mQH_2_O: MilliQ water
MTG: Middle temporal gyrus
PBS-T: PBS with Triton X-100
PBS: Phosphate-buffered saline
PMD: Post-mortem delay
PolyQ: Polyglutamine
PRD: Proline-rich domain
TMA: Tissue microarray

