## Supplementary Figure 1 for "N-terminal mutant Huntingtin deposition correlates with CAG repeat length and disease onset, but not neuronal loss in Huntington’s disease"

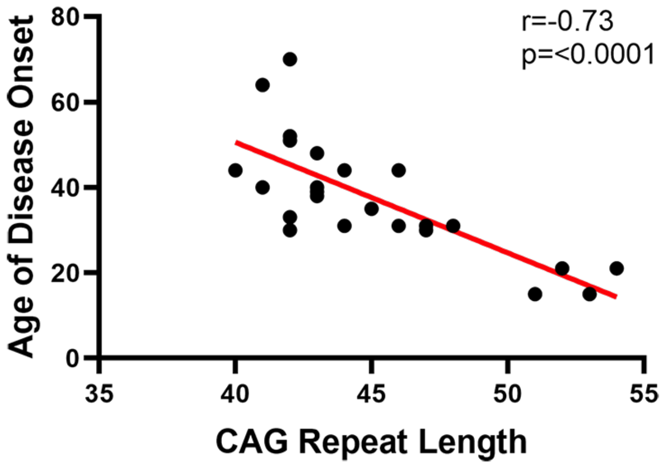


**Supplementary Figure 1.** Correlation between CAG repeat length and age of disease onset within the HD TMA cohort (n=28). A strong negative correlation was found between the CAG repeat length and age of disease onset (r=-0.73; p=<0.0001).
