## Supplementary Table 1 for "N-terminal mutant Huntingtin deposition correlates with CAG repeat length and disease onset, but not neuronal loss in Huntington’s disease"

**Supplementary Table 1** Summary of successful and unsuccessful anti-Htt antibodies applied to paraffin-embedded HD and control human brain tissue using immunohistochemistry. Table includes all IHC conditions tested for each antibody: antibody concentration (up to four different concentrations were tested), antigen retrieval buffer, and DAB incubation time. All antibodies were treated with formic acid in addition to the antigen retrieval buffer listed. The optimal IHC conditions for the successful antibodies (highlighted in green, above dashed line) which displayed immunoreactivity are also included. The antibodies denoted with an asterisk immunolabeled Htt aggregates. IHC = immunohistochemistry; DAB = 3,3'-diaminobenzidine; TMA = tissue microarray; HD = Huntington's disease; Htt = Huntingtin.

| **Marker** | **IHC optimization conditions tested in whole sections** | | | **Optimal conditions for TMA IHC:** Concentration; Antigen retrieval buffer; DAB time (minutes) |
|---|---|---|---|---|
| | Concentration | Antigen retrieval buffer (plus Formic Acid) | DAB time (minutes) | |
| MAB5374* | 1:100 | Sodium-Citrate pH 6.0; Tris-EDTA pH 9.0 | 15 | 1:100; Sodium-Citrate pH 6.0; 15 |
| MW1* | 1:100; 1:500; 1:1000; 1:2500 | Tris-EDTA pH 9.0 | 7; 10 | 1:1000; Tris-EDTA pH 9.0; 7 |
| EPR5526* | 1:100; 1:500; 1:1000: 1:2500; 1:5000 | Sodium-Citrate pH 6.0; Tris-EDTA pH 9.0 | 5; 7; 10 | 1:5000; Sodium-Citrate pH 6.0; 7 |
| MAB5492 | 1:100, 1:500, 1:1000 | Tris-EDTA pH 9.0 | 10 | 1:500; Tris-EDTA pH 9.0; 10 |
| MW8 | 1:100; 1:500; 1:1000 | Sodium-Citrate pH 6.0; Tris-EDTA pH 9.0 | 5; 10 | 1:500; Tris-EDTA pH 9.0; 5 |
| 2B7 | 1:500; 1:1000; 1:2000 | Sodium-Citrate pH 6.0; Tris-EDTA pH 9.0 | 5 | 1:2000; Sodium-Citrate pH 6.0; 5 |
| 2E10 | 1:500; 1:1000 | Sodium-Citrate pH 6.0; Tris-EDTA pH 9.0 | 3; 5 | - |
| 4C9 | 1:500; 1:1000 | Sodium-Citrate pH 6.0; Tris-EDTA pH 9.0 | 5 | - |
| D7F7 | 1:250; 1:500 | Sodium-Citrate pH 6.0; Tris-EDTA pH 9.0 | 10 | - |
| MAB2166 | 1:20; 1:100; 1:500 | Sodium-Citrate pH 6.0; Tris-EDTA pH 9.0 | 10 | - |
| MAB2168 | 1:500; 1:750; 1:1000 | Sodium-Citrate pH 6.0; Tris-EDTA pH 9.0 | 5; 10 | - |
| MAB5490 | 1:100; 1:200; 1:250; 1:500 | Sodium-Citrate pH 6.0; Tris-EDTA pH 9.0 | 10; 15 | - |
